# The hyaluronan-binding activity of aggrecan is important, but not essential, for its specific insertion into perineuronal nets

**DOI:** 10.1101/2024.11.25.625086

**Authors:** Matthew Y. Otsuka, Leslie B. Essel, Ashis Sinha, Gabrielle Nickerson, Seth M. Mejia, Russell T. Matthews, Samuel Bouyain

## Abstract

Aggrecan (ACAN) is a large, secreted chondroitin sulfate proteoglycan that includes three globular regions named G1, G2, G3, and is decorated with multiple glycosaminoglycan attachments between its G2 and G3 domains. The N-terminal G1 region interacts with the glycosaminoglycan hyaluronan (HA), which is an essential component of the vertebrate extracellular matrix. In the central nervous system, ACAN is found in perineuronal nets (PNNs), honeycomb-like structures that are enriched on parvalbumin-positive neurons in specific neural circuits. PNNs regulate the plasticity of the central nervous system, and it is believed that association between ACAN and HA is a foundational event in the assembly of these reticular structures. Here, we report the co-crystal structure of the G1 region of ACAN in the absence and presence of an HA decasaccharide and analyze the importance of the HA-binding activity of ACAN for its integration into PNNs. We demonstrate that the single immunoglobulin domain and the two Link modules that comprise the G1 region form a single structural unit, and that HA is clamped inside a groove that spans the length of the tandem Link domains. Introduction of point mutations in the glycosaminoglycan-binding site eliminates HA-binding activity in ACAN, but, surprisingly, only decreases the integration of ACAN into PNNs. Thus, these results suggest that the HA-binding activity of ACAN is important for its recruitment to PNNs, but it does not appear to be essential.

## INTRODUCTION

The glycosaminoglycan hyaluronan (HA) is an abundant component of vertebrate extracellular matrices (ECMs) (1). Unlike other linear, negatively charged, polysaccharides such as heparan sulfate or chondroitin sulfate, HA is neither sulfated nor covalently attached to proteins secreted in the extracellular environment or found at the cell membrane. Instead, HA is assembled from the monosaccharides *N*-acetylglucosamine (GlcNAc) and glucuronic acid (GlcUA) found in the cytoplasm by hyaluronan synthases. Humans express three hyaluronan synthases named HAS1-3, and these channel-like proteins embedded in the plasma membrane translocate HA chains composed of repeating units of the disaccharide β-1,3-GlcNAc-β-1,4- GlcUA chains into the extracellular environment. Extruded HA chains may exceed several megadaltons in molecular weight (2). The simple structure of HA perhaps belies the manifold essential functions it has throughout the lifetime of an individual, from mediating morphogenesis in embryos to scaffolding of the ECM of numerous tissues such as cartilage and brain during adulthood (3–6).

The diverse physiological functions of HA are linked on the one hand to its unique water-retention and physicochemical properties and on the other hand to its interactions with cell surface receptors or ECM proteins. Several of these HA-binding proteins include one or more ∼100 amino acids repeats called Link modules that were initially identified in two ECM proteins called aggrecan (ACAN) and hyaluronan and proteoglycan link protein 1 (HAPLN1) found in porcine cartilage (7, 8). *Hapln1-* or *Acan-*knockout mice die soon after birth and present defects in the development of cartilage and bone, underlying the importance of ACAN and HAPLN1 in normal mammalian development (9–11). ACAN is a large, secreted glycoprotein that spans more than 2,500 amino acids in humans (12). Its N-terminus includes a globular region named G1 that shares ∼ 39% amino acid identity with HAPLN1, and is composed of an immunoglobulin (Ig) domain and two link modules (Fig. 1A) (13). The G1 region of ACAN as well as HAPLN1 bind to HA through their Link modules (14, 15) and biophysical analyses suggest that the N-terminus of ACAN forms a ternary complex with HAPLN1 and HA (16), possibly explaining why *Hapln1-* or *Acan-*knockout mice exhibit similar defects in cartilage.

**Figure 1.**
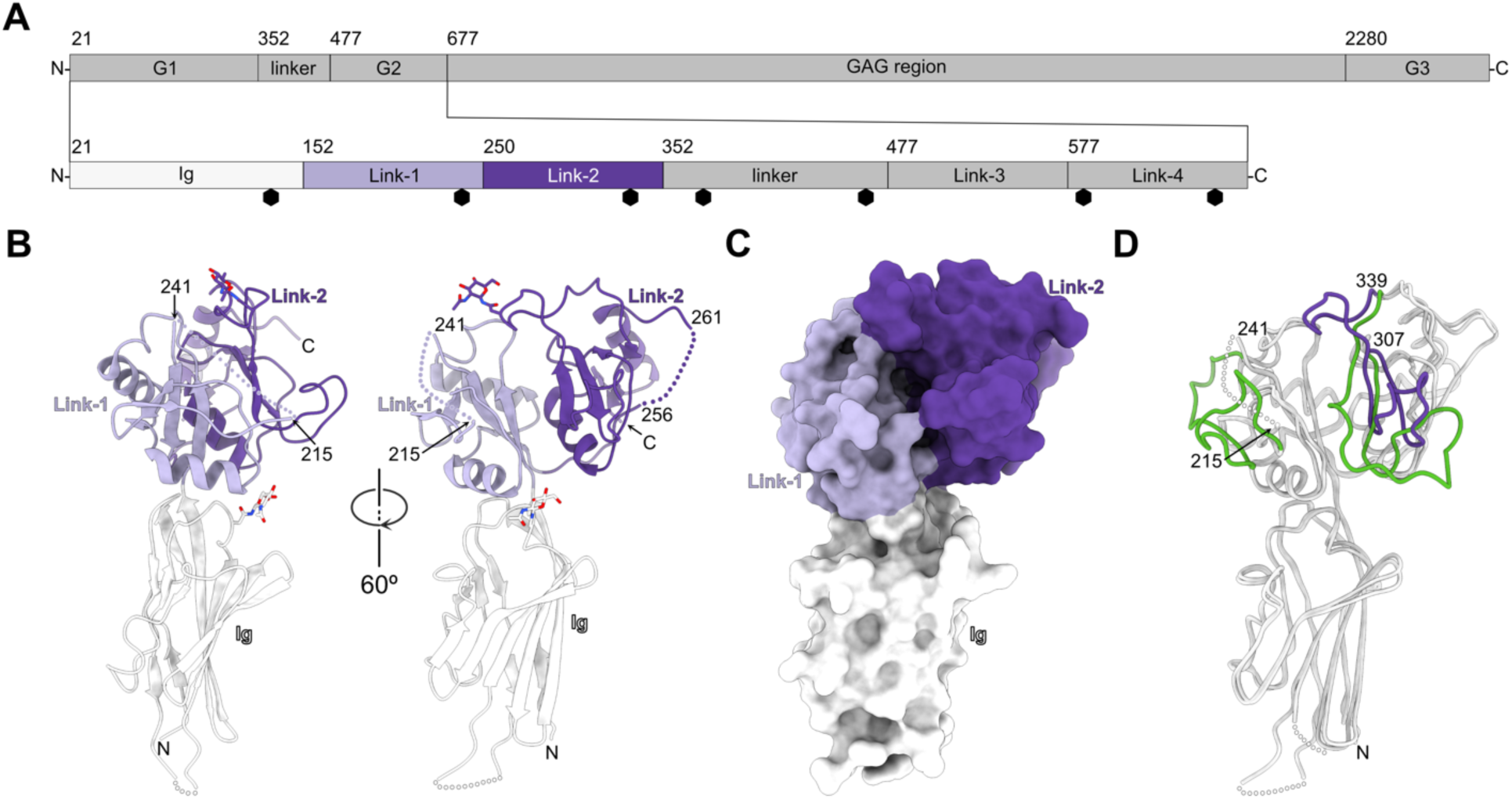
The crystal structure of the G1 domain of ACAN. A. Overview of the domain organization of human ACAN. ACAN includes three globular regions named G1, G2, and G3. A large ∼1,600 amino acid segment between the G2 and G3 domains includes attachment sites for covalent modification with sulfated glycosaminoglycans (GAGs), such as chondroitin sulfate. The G1-G2 segment includes an N-terminal immunoglobulin (Ig) domain and four Link modules. The Ig, Link-1, and Link-2 domains form the G1 region while the Link-3 and Link-4 domains form the G2 region. The Ig, Link-1, and Link-2 are colored white, lilac, and violet, respectively. This color coding is used in panels B and C below. Amino acid numbering corresponds to human ACAN (Uniprot ID# P16112). Black hexagons denote predicted N-linked glycosylation sites. B. The G1 region of human ACAN is shown in a ribbon diagram in two distinct orientations related by a 60° rotation. The Ig, Link-1, and Link-2 domains in chain B of ACAN are colored white, lilac, and violet, respectively. The letters N and C indicate the N- and C- termini, respectively. Asparagine-linked N-acetylglucosamine residues are shown as sticks along with the asparagine side chain. Three disordered regions in the Ig, Link-1, and Link-2 domains are shown as dotted lines. C. Surface representation of the G1 domain of ACAN. Domains are colored as described in panel B. ACAN(G1) is shown in the same orientation as the on right view in panel B. D. Overlay of the structures of the two chains of ACAN(G1) in the asymmetric unit of ACAN(G1) crystals showing distinct conformations for chains A and B. The two chains are shown as coils and colored white. Residues 216-240 are not visible in chain B but are well ordered in chain A and shown in green. Residues 307-339 in chain A are colored green while the same region in chain B is colored violet. ACAN, aggrecan; GAG, glycosaminoglycan; Ig, immunoglobulin;

The physiological functions of ACAN and HAPLN1 are not limited to bone and cartilage, however. In the central nervous system (CNS) these two proteins are essential components, along with HA, of a condensed form of ECM called perineuronal nets (PNNs) that form around a specific subset of neurons (17–20). Broadly, the role of PNNs is to limit neuroplasticity, the ability of the brain to alter its networks as a result of neural activity (21). PNNs are reticular structures that form around the soma and dendrites of subpopulations of neurons throughout the CNS. The presence of nets generally correlates with the expression of the Ca^2+^-binding protein parvalbumin and they encase neurons concomitantly with the closing of a period of intense neural remodeling in response to experience called the critical period (22–25). Although the exact molecular composition of PNNs is thought to vary with the identity of the neural tissue in which they form (26), HA, ACAN, HAPLN1, the matricellular protein tenascin-R (TNR), and the secreted proteoglycan phosphacan appear to be essential for the formation of nets (17, 19, 27, 28). Accordingly, genetic ablation of *Acan*, *Hapln1*, or *Tnr* restores plasticity in mice (17, 19, 29). Crucially, although ACAN is expressed ubiquitously outside of the nervous system, it appears to be found almost exclusively in PNNs in neural tissues (30, 31).

Since (i) ACAN is an essential component of PNNs (19) and (ii) the expression of hyaluronan synthase 3 (HAS3) in human embryonic kidney cells induces the capture of ACAN in a condensed matrix surrounding these cells (18), it was of interest to determine the extent to which the integration of ACAN into PNNs depends on its physical interaction with HA. In this report, we provide the crystal structures of the G1 region of human ACAN in the absence and presence of an HA oligosaccharide. Mutations of four residues in the glycosaminoglycan-binding site found in the tandem Link domains of ACAN eliminated its HA-binding activity in BioLayer Interferometry assays. However, in the context of neuronal cultures, these mutations impaired, but did not eliminate the integration of ACAN into PNNs. Overall, these results suggest that the HA-binding activity is not obligatory for its recruitment into this specialized form of neural ECM, suggesting that additional interactions, perhaps between ACAN and another protein, are also important for the assembly of PNNs.

## RESULTS

### The Ig and the two Link modules in the G1 domain of ACAN form a single structural unit

ACAN includes three globular regions named G1, G2, and G3 with a large insert for attachment of sulfated glycosaminoglycans between the G2 and G3 domains (Fig. 1A). The G1- G2 region includes four Link domains arranged in two pairs: the N-terminal Ig domain and the Link-1 and Link-2 pair form the G1 domain that binds to HA (14) while the Link-3 and Link-4 modules form a second pair that corresponds to the G2 region. The crystal structure of the G1 region of ACAN was determined as a first step to investigate the association of ACAN and HA. The structure of deglycosylated ACAN(G1) was solved by molecular replacement using a model generated by AlphaFold (32) and refined to 3.50 Å (Rwork / Rfree = 0.247 / 0.284, Table 1, Fig. 1B- C and Fig. S1). There are two molecules of ACAN in the crystal asymmetric unit. The N-terminal Ig domain of ACAN comprises amino acids L29 – K151, while the Link-1 and Link-2 modules span G152 – E249 and G250 – T350, respectively (Fig. 1A). Overall, these three discrete domains form a single structural unit that resembles an inverted letter “L” with dimensions of ∼ 60 x 50 x 20 Å. The interface between the Ig domain and Link-1 buries 430 Å^2^, while the interface between Link-1 and Link-2 buries 682 Å^2^ (Fig. 1B, C). The Ig domain does not contact Link-2.

**Table 1 -.** Data collection and refinement statistics. Values in parentheses apply to the high-resolution shell

|  | ACAN(G1) | ACAN(G1)-HA<sub>10</sub> complex |
| --- | --- | --- |
| Data collection |  |  |
| Beamline | APS 22-ID | APS 22-ID |
| Wavelength (Å) | 1.00 | 1.00 |
| Number of unique reflections | 20,877 (4,880) | 29,486 (2,940) |
| Resolution (Å) | 85.91 – 3.50 | 50.00 – 2.58 |
| Space group | P4<sub>1</sub>2<sub>1</sub>2 | P2<sub>1</sub> |
| Unit cell |  |  |
| a, b, c (Å) | 121.49, 121.49, 213.37 | 73.07, 64.94, 112.21 |
| α, β, γ (°) | 90.00, 90.00, 90.00 | 90.00, 109.33, 90.00 |
| R<sub>merge</sub> | 0.125 (1.324) | 0.163 (0.542) |
| R<sub>pim</sub> | 0.044 (0.479) | 0.104 (0.361) |
| Completeness (%) | 100.00 (100.00) | 96.3 (96.9) |
| Redundancy | 8.9 (9.3) | 3.2 (2.8) |
| I/σI | 7.5 (2.4) | 6.1 (1.5) |
| CC<sub>1/2</sub> | 0.995 (0.683) | 0.968 (0.731) |
| Refinement |  |  |
| PDB code | 9DFT | 9DFF |
| Number of protein chains in asymmetric unit | 2 | 2 |
| Resolution (Å) | 61.38 – 3.50 | 41.03 – 2.59 |
| Reflections (Test) | 20,824 (2,084) | 27,805 (1,884) |
| R<sub>work</sub> / R<sub>free</sub> | 0.198 / 0.245 | 0.194 / 0.238 |
| Atoms modeled | 4,758 | 5,327 |
| Protein | 4,678 | 4,810 |
| Ligand | 80 | 344 |
| Water | - | 173 |
| R.m.s deviations |  |  |
| Ideal bonds (Å) | 0.002 | 0.005 |
| Ideal angles (°) | 0.507 | 0.770 |
| Average B factors (Å<sup>2</sup>) | 167.78 | 45.80 |
| Protein | 166.96 | 45.33 |
| Ligand | 216.13 | 56.57 |
| Water | - | 37.22 |
| Ramachandran statistics |  |  |
| Favored (%) | 95.37 | 96.13 |
| Allowed (%) | 4.46 | 3.53 |
| Outlier (%) | 0.17 | 0.34 |
| Rotamer outlier (%) | 0.20 | 0.00 |

The two protomers in the asymmetric unit superimpose with a root mean square deviation (RMSD) of 1.25 Å over 251 Cα pairs. However, examination of the two protein chains suggests that there are significant differences in the arrangement of loop regions in the Link domains between the two protomers (Fig. 1D). Specifically, although continuous electron density in chain A made it possible to build the entire Link tandem repeats (Fig. S1A), two segments in the Link domains of chain B could not be built because of missing or uninterpretable electron density (Fig. 1B). These segments include amino acids 216 – 240 and 257 – 260. Furthermore, the conformation of region 307 – 339 in chain A differs significantly from the one adopted in chain B (Fig. 1D). The conformation of this segment in chain A may be an artifact of crystallization because the single GlcNAc attached at N333 is wedged between the Link-1 and Link-2 domains (Fig. S1A, C). This monosaccharide residue remained attached to ACAN(G1) after treatment with endoglycosidase H prior to crystallization. In a non-deglycosylated protein, however, the bulk of the asparagine-linked complex carbohydrate would be likely to prevent such positioning, suggesting that the conformation of the region 307 – 339 in chain A may not reflect its conformation in solution. In addition, amino acids 312 – 330 in chain A protrude away from the Link-2 domain to participate in crystal contact with two symmetry-related mates (Fig. S1A). As such, it appears likely that the conformation adopted by chain B is more physiologically relevant than the one adopted by chain A.

### The two Link domains assemble a contiguous binding site that accommodates a single HA decamer

Previous work has indicated that ACAN may associate with HA oligosaccharide that comprises at least ten monosaccharides (16). Thus, the G1 domain of ACAN was crystallized in the presence of a fivefold molar excess of a HA decamer (HA_10_) to obtain atomic level information between ACAN and HA. The crystal structure of the ACAN(G1)-HA_10_ complex was solved by molecular replacement and refined to 2.59 Å (Rwork / Rfree = 0.206 / 0.240, Table 1, Fig. 2, Fig. S2). There are two molecules of ACAN in the asymmetric unit that superimpose with a RMSD of 0.18 Å over 301 Cα positions. The electron density for the bound HA_10_ is well-defined for all the sugar residues in the oligosaccharide (Fig. 2B and Fig. S2). The oligosaccharide buries 820 Å^2^ of surface area and fits within a groove that spans both Link domains in ACAN, while the Ig domain does not make any contact with HA (Fig. 2A). Comparison of the bound and free form of ACAN indicates that the segment comprising residues 217 – 241 that was disordered in HA-free ACAN is now ordered and makes substantial contact with the oligosaccharide. Concomitantly, HA_10_ adopts an almost linear conformation that differs from the structure of free HA obtained from powdered diffraction (Fig. S3). This change is consistent with the alteration of the HA structure reported upon binding to ACAN(G1) (33). Finally, comparing the HA-binding site in ACAN and in Cd44, the other available complex between a Link module and a fragment of HA (34), shows that the sugar-binding grooves in Link-1 and Link-2 both match the HA-binding site in Cd44 (Fig. S4A). However, a loop extends from Link-2 to bind to HA between the two Link domains, giving the appearance that the oligosaccharide is pinched at the interface between Link1 and Link2. This loop is absent in Cd44 and corresponds to an insertion of 8 amino acids between residue 109 and 110 of Cd44 (Fig. S4B).

**Figure 2.**
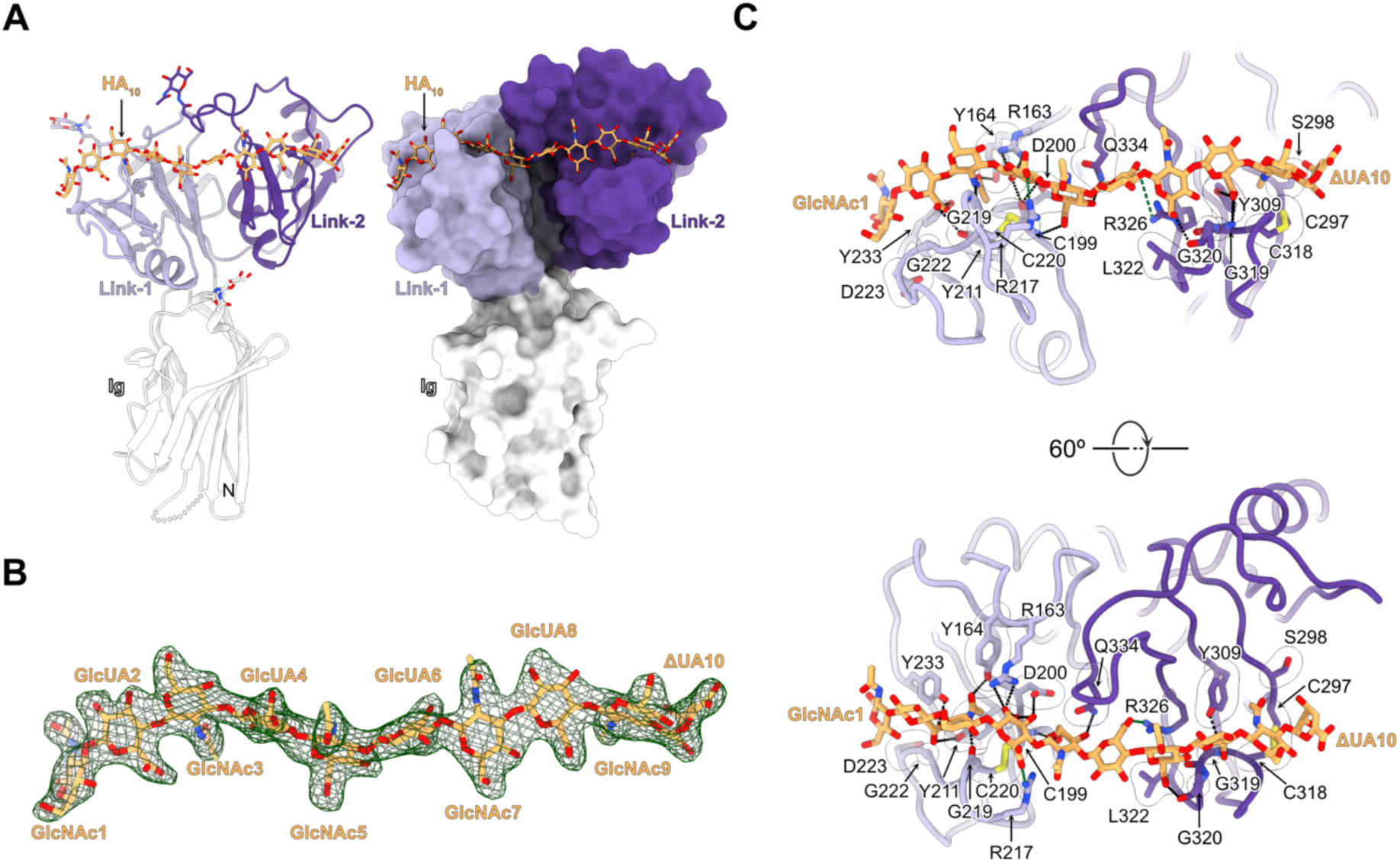
Crystal structure of the complex between ACAN and a HA oligosaccharide. A. The G1 domain of ACAN bound to HA_10_ is shown as a ribbon diagram on the left and in surface representation on the right. The bound HA decasaccharide is shown as sticks in both views and colored orange. The Ig, Link-1, and Link-2 domains in chain B of ACAN are colored white, lilac, and violet, respectively. The letters N and C indicate the N- and C-termini, respectively. Asparagine-linked N-acetylglucosamine residues are shown as sticks along with the asparagine side chain. B. mFo-DFc electron density map for the bound HA_10_ calculated by omitting the oligosaccharide from the model is contoured at 3 σ and shown as a green mesh. The oligosaccharide is in the same orientation as the one shown in panel A. Residues in the HA oligosaccharide are colored orange. The terminal residue, 4-deoxy-alpha-L-threo- hex-4-enopyranuronic acid, created during the preparation of the HA oligosaccharide, is labeled ΔUA. C. Detailed view of the interactions between HA_10_ and residues in the Link-1 and Link-2 of ACAN(G1). The view on top is in the same orientation as the view shown in panel A. The bound HA is shown in stick representation and colored orange. Link-1 and Link-2 are shown as coils. Contacting protein residues are represented as sticks, while transparent surfaces denote residues involved in van der Waals or packing interactions. Potential hydrogen bonds are represented as black dashed lines between interacting atoms while salt bridge interactions involving R217 and R326 are shown as green dashed lines. Interacting atoms of side chains or main chain atoms in ACAN are displayed only if they participate in the interactions with HA_10_. Only the first and tenth residue of HA_10_ are labeled for the sake of clarity. Residues in Link-1 are colored lilac, while residues in Link- 2 are colored violet. ACAN, aggrecan; GlcNAc, *N*-acetylglucosamine; GlcUA, beta-D-glucopyranuronic acid; HA, hyaluronan; Ig, immunoglobulin; ΔUA, 4-deoxy-alpha-L-threo-hex-4-enopyranuronic acid.

Overall, the ten monosaccharide residues in the crystallized HA oligosaccharide contact eighteen amino acids in ACAN through van der Waals or hydrogen bonding interactions (Fig. 2C). All but three of these eighteen HA-contacting residues appear well conserved among vertebrates ACAN proteins (Fig. S5). In contrast to the interface between Cd44 and HA, only one aliphatic amino acid, L322, contacts the bound HA (Fig. S4B). Instead, van der Waals contacts occur between the side chains of C199, D200, Y211, D223, C297, S298, Y309, L322, and Q334 on the one hand and GlcNAc1, GlcNAc3, GlucA4, GlucA6, GlcNAc7, GlucA8, GlcNAc9, and the terminal ΔUA10 on the other hand. Furthermore, sugar residues GlucA2- GlcA8 form an extensive network of eighteen hydrogen bonds with ACAN. These include the main chain atoms of C199, G219, G319, and G320 forming seven hydrogen bonding interactions with GlcNAc3, GlucA4, GlcNAc7 and GlucA8. In addition, the side chain atoms of R163, Y164, D200, Y211, R217, Y233 in Link1 contact GlucA2 to GlcNAc5 while the side chains of Y309, R326, and Q334 form hydrogen bonds with GlcNAc5 to GlucA8. Surprisingly, there are only two salt bridge interactions between HA and ACAN residues in spite of the five negative charges found in HA. These interactions involve the guanidinium groups of R217 and R326, which contact the carboxylate groups of GlucA4 and GlucA6 at the center of the bound decasaccharide (Fig. 2C).

### Mutations in the Link-1 and Link-2 domains of ACAN impair binding to HA

A binding assay using bio-layer interferometry (BLI) was designed to quantify the interaction between HA and ACAN(G1) as well as validate the structural results described above. In these experiments, the binding of increasing concentrations of ACAN(G1) to a 20-kDa fragment of biotinylated HA immobilized on a streptavidin biosensor was measured. The traces were fit to a 1:1 model and the apparent binding affinity (K_D_) was calculated from the rates of association (k_on_) and dissociation (k_off_) (Fig. 3A). In these experiments, ACAN binds to immobilized HA with an affinity of 234 nM, which is consistent with the value of 226 nM reported in an earlier study using surface plasmon resonance (14).

**Figure 3.**
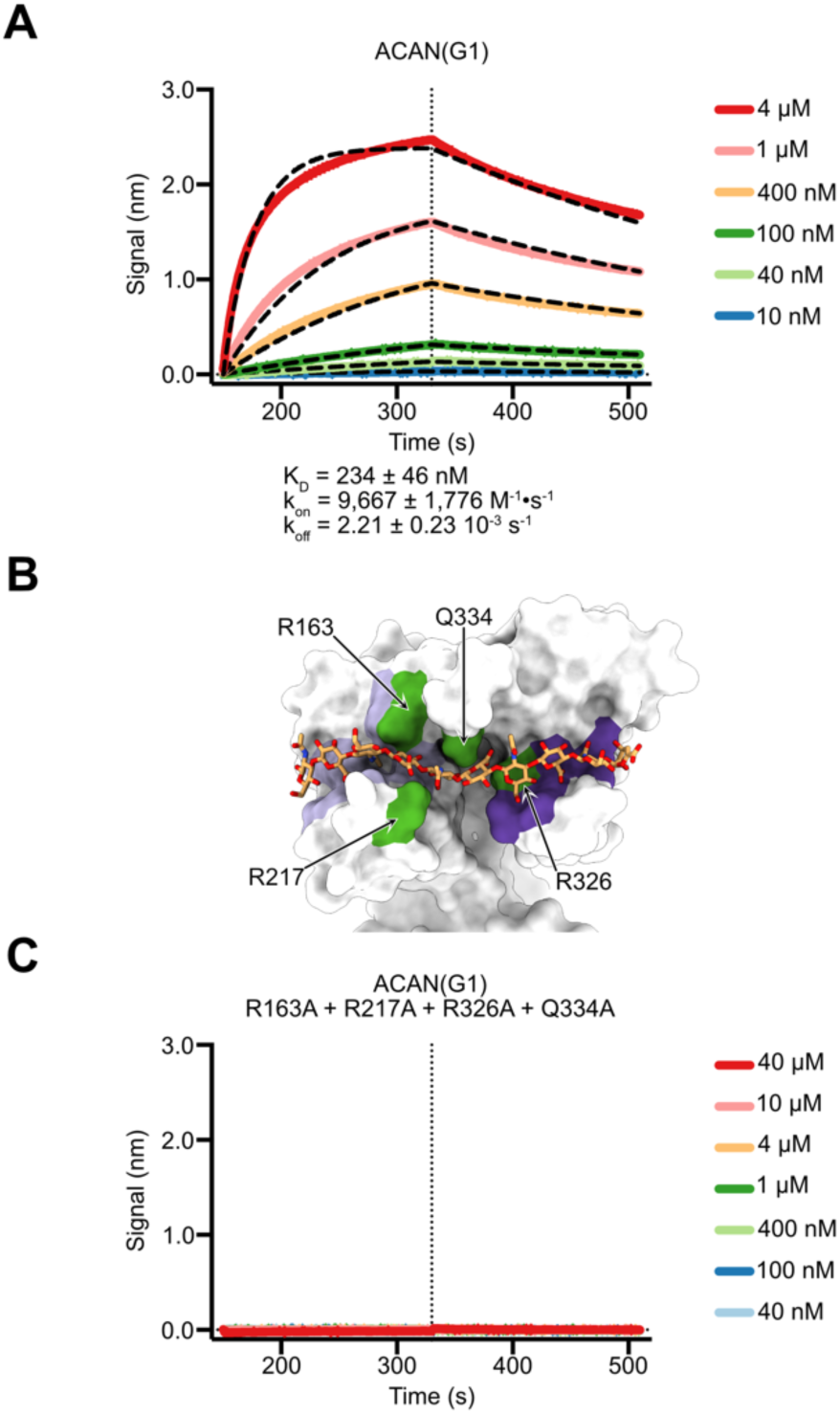
Characterization of ACAN(G1) binding to immobilized HA by biolayer interferometry. A. A 20-kDa fragment of HA including a single biotin residue was immobilized onto streptavidin sensors, and titrated against 4 μM, 1 μM, 400 nM, 100 nM, 40 nM, and 10 nM of the G1 domain of human ACAN. Representative association and dissociation curves are shown here along with a vertical dashed line that indicates the start of the dissociation phase. Raw binding data are shown in distinct colors and were analyzed using a 1:1 binding model. Curves obtained from fitting are shown as black dashed lines. Values for the affinity constant K_D_, the rate of association k_on_, and the rate of dissociation k_off_ are reported as averages ± SDs from eleven independent experiments using three distinct biological replicates. Additional information about individual experiments used in affinity calculations are listed in Table S1. B. The tandem Link modules in ACAN(G1) are shown as a white surface along with the bound HA shown as orange sticks. HA-binding residues in the Link-1 module are colored lilac, while HA-binding residues in Link-2 are colored violet. The positions of residues that are changed alanine in the HA-binding mutant of ACAN(G1) are colored green. This view is in the same orientation as the one shown in the bottom view of Fig. 2C. C. A 20-kDa fragment of HA including a single biotin residue was immobilized onto streptavidin sensors, and titrated against 40 µM, 10 µM, 4 μM, 1 μM, 400 nM, 100 nM, and 40 nM of the G1 domain of human ACAN including mutations to alanine at positions R163, R217, R326, and Q334. Representative association and dissociation curves are shown here along with a vertical dashed line that indicates the start of the dissociation phase. The raw binding data did not show any interaction between immobilized HA and this variant of ACAN(G1). ACAN, aggrecan; HA, hyaluronan.

The binding assay validated, four mutations to alanine were introduced at the interface between ACAN(G1) and the bound HA_10_ to confirm that the contacts observed in the crystals reflect interactions in solution. Residues R163, R217, R326, and Q334 were selected because they interact with the trisaccharide GlucA4-GlcNAc5-GlucA6 located centrally in HA_10_ and appear strictly conserved in selected vertebrate species (Fig. 2C, 3B, S5). The mutant protein behaved comparably with wild-type ACAN(G1) during purification, specifically size exclusion chromatography, suggesting that the introduction of the mutations did not introduce gross structural alterations (data not shown). In BLI assays, the mutant ACAN(G1) did not associate with immobilized HA (Fig. 3C). Overall, these results indicate that the protein-HA interface identified in the complex structure is relevant to interactions in solution.

**Figure 4.**
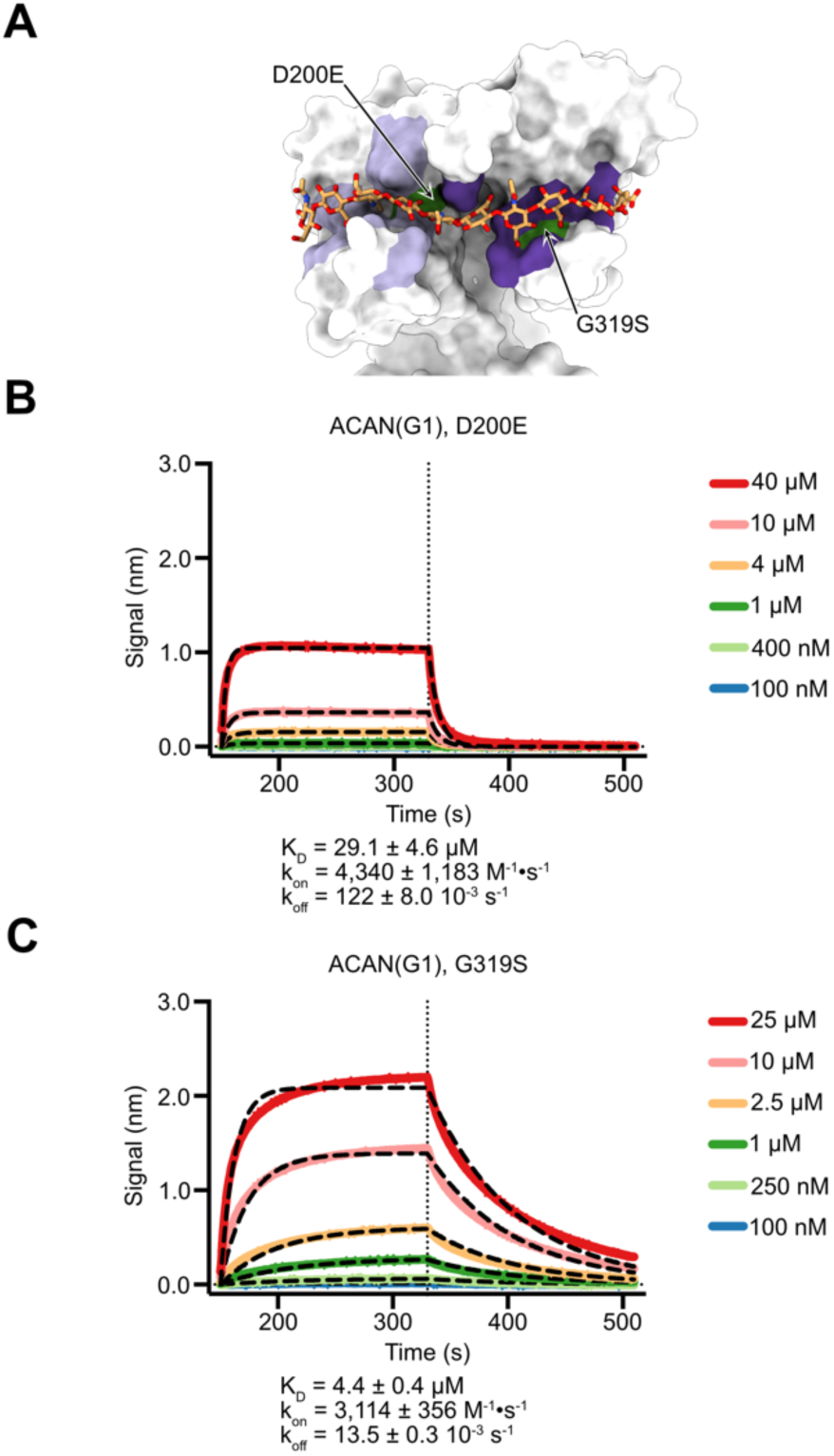
Characterization of missense mutations found in the HA-binding site of ACAN(G1) associated with osteochondritis dissecans. A. The tandem Link modules in ACAN(G1) are shown as a white surface along with the bound HA shown as orange sticks. HA-binding residues in the Link-1 module are colored lilac, while HA-binding residues in Link-2 are colored violet. The positions of the two missense mutations in the HA-binding site associated with osteochondritis dissecans are colored green. This view is in the same orientation as the one shown in the bottom view of Fig. 2C. B. A 20-kDa fragment of HA including a single biotin residue was immobilized onto streptavidin sensors, and titrated against 40 µM, 10 µM, 4 μM, 1 μM, 400 nM, 100 nM of human ACAN(G1) including the change D200E. Representative association and dissociation curves are shown here along with a vertical dashed line that indicates the start of the dissociation phase. Raw binding data are shown in distinct colors and were analyzed using a 1:1 binding model. Curves obtained from fitting are shown as black dashed lines. Values for the affinity constant K_D_, the rate of association k_on_, and the rate of dissociation k_off_ are reported as averages ± SDs from five independent experiments using two distinct biological replicates. Additional information about individual experiments used in affinity calculations are listed in Table S1. C. A 20-kDa fragment of HA including a single biotin residue was immobilized onto streptavidin sensors, and titrated against 25 µM, 10 µM, 2.5 μM, 1 μM, 250 nM, 100 nM of the G1 domain of human ACAN including the G319S mutation. Representative association and dissociation curves are shown here along with a vertical dashed line that indicates the start of the dissociation phase. Raw binding data are shown in distinct colors and were analyzed using a 1:1 binding model. Curves obtained from fitting are shown as black dashed lines. Values for the affinity constant K_D_, the rate of association k_on_, and the rate of dissociation k_off_ are reported as averages ± SDs from five independent experiments using two distinct biological replicates. Additional information about individual experiments used in affinity calculations are listed in Table S1. ACAN, aggrecan; HA, hyaluronan.

**Figure 5.**
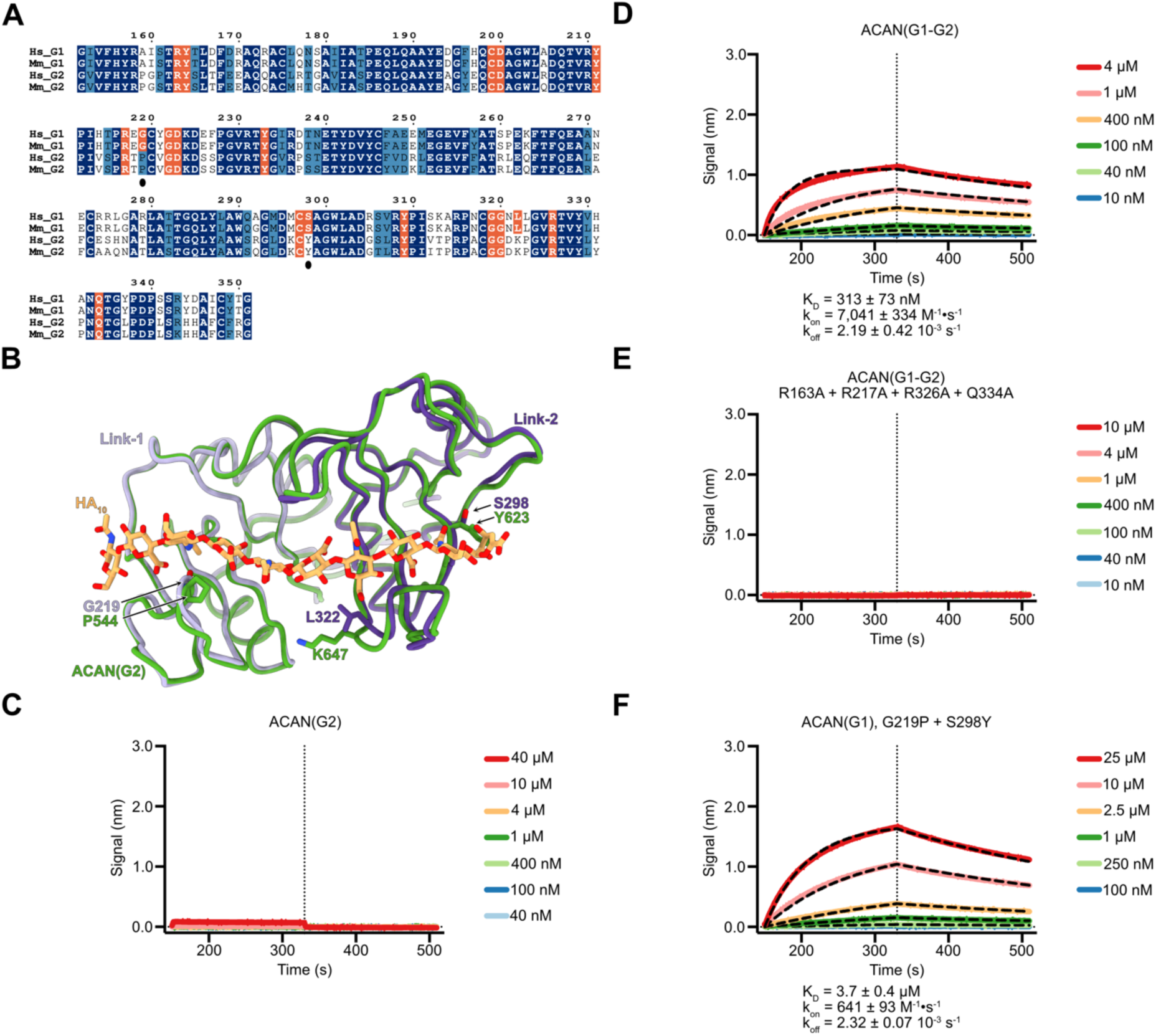
The tandem Link modules that make up the G2 domain of ACAN do not bind HA. A. Amino acid conservation between the tandem Link modules found in the G1 and G2 regions of human and mouse ACAN. The numbering corresponds to human ACAN(G1). Identical residues are shaded in navy while similar residues are shaded in blue. Residues that interact with HA in human ACAN are shaded orange. They are highlighted in the same color in other sequences to indicate residue conservation. Black dots under G219 and S298 indicate HA-binding residues that are distinct between the G1 and G2 domains and are mutated in the experiments shown in panel F. B. Overlay of the structures of the tandem Link modules in ACAN(G1) and in ACAN(G2). The model for ACAN(G2) was generated using AlphaFold3. Proteins are shown in coil representation while the bound oligosaccharide is shown as orange sticks. The Link-1 and Link-2 domains in ACAN(G1) are colored lilac, and violet, respectively, while the Link-3 and Link-4 modules in ACAN(G2) are colored green. The three HA-binding residues in ACAN(G1) that are distinct in ACAN(G2) are shown as sticks and colored lilac or violet. The corresponding residues in ACAN(G2) are also shown as sticks and colored green. C. A 20-kDa fragment of HA including a single biotin residue was immobilized onto streptavidin sensors, and titrated against 40 µM, 10 µM, 4 μM, 1 μM, 400 nM, 100 nM, and 40 nM of the G2 domain of human ACAN. Representative association and dissociation curves are shown here along with a vertical dashed line that indicates the start of the dissociation phase. The raw binding data did not show any interaction between immobilized HA and ACAN(G2). D. A 20-kDa fragment of HA including a single biotin residue was immobilized onto streptavidin sensors, and titrated against 4 μM, 1 μM, 400 nM, 100 nM, 40 nM, and 10 nM of the G1-G2 region of human ACAN. Representative association and dissociation curves are shown here along with a vertical dashed line that indicates the start of the dissociation phase. Raw binding data are shown in distinct colors and were analyzed using a 1:1 binding model. Curves obtained from fitting are shown as black dashed lines. Values for the affinity constant K_D_, the rate of association k_on_, and the rate of dissociation k_off_ are reported as averages ± SDs from seven independent experiments using three distinct biological replicates. Additional information about individual experiments used in affinity calculations are listed in Table S1. E. A 20-kDa fragment of HA including a single biotin residue was immobilized onto streptavidin sensors, and titrated against 10 µM, 4 μM, 1 μM, 400 nM, 100 nM, 40 nM, and 10 nM of the G1-G2 region of human ACAN including mutations to alanine at positions R163, R217, R326, and Q334. Representative association and dissociation curves are shown here along with a vertical dashed line that indicates the start of the dissociation phase. The raw binding data did not show any interaction between immobilized HA and this variant of ACAN(G1-G2). F. A 20-kDa fragment of HA including a single biotin residue was immobilized onto streptavidin sensors, and titrated against 25 µM, 10 µM, 2.5 μM, 1 μM, 250 nM, 100 nM of human ACAN(G1) including the changes G219P and S298Y. Representative association and dissociation curves are shown here along with a vertical dashed line that indicates the start of the dissociation phase. Raw binding data are shown in distinct colors and were analyzed using a 1:1 binding model. Curves obtained from fitting are shown as black dashed lines. Values for the affinity constant K_D_, the rate of association k_on_, and the rate of dissociation k_off_ are reported as averages ± SDs from four independent experiments using two distinct biological replicates. Additional information about individual experiments used in affinity calculations are listed in Table S1. ACAN, aggrecan; HA, hyaluronan. Hs, *Homo sapiens*; Mm, *Mus musculus*.

Mutations in ACAN have been linked to several human diseases that affect bone and/or cartilage (35, 36). Thus, it was of interest to examine whether reported mutations map to the HA-binding site and what their effect on HA binding might be. We searched the ClinVar database for missense mutation in ACAN and identified two changes in residues that contact HA in the crystal structure (Fig. 2C) (37). These residues also appear conserved in the G1 domains of selected vertebrate species (Fig. S5). These two mutations, a change from aspartate to glutamate at position 200 (D200E) in Link-1 and a change from glycine to serine at position 319 (G319S) in Link-2 are associated with osteochondritis dissecans, a condition that affects bone and cartilage (Fig. 4A) (38). ACAN(G1) proteins including these changes were expressed and their HA-binding properties assessed using BLI (Fig. 4). Introduction of the amino acid changes did not appear to introduce any gross alteration of the G1 domain as the mutant proteins could be purified to homogeneity from conditioned media as easily as the wild type protein.

Although changes from aspartate to glutamate are often considered benign, introducing this mutation at position 200 decreased the affinity of ACAN(G1) for HA more than a hundredfold (29.1 µM versus 234 nM, Fig. 4B). The carboxylate group of D200 forms a hydrogen bond with a hydroxyl group in GlucA4 and introduction of an additional methyl group in the side chain might put the carboxylate group too far for such an interaction. In addition, D200 lies deep in the HA-binding crevice so that elongation of the side chain after the change to glutamate might push the HA chain away from the binding site because of a reduction in the size of the HA-binding groove. The change at position 319 from a glycine to a serine does not impact the affinity for HA to the same extent as D200E, but the dissociation constant is still decreased almost twentyfold (4.4 µM versus 234 nM). The main chain nitrogen atom of G319 forms a hydrogen bond with residue GlucA8. Because this interaction would not be expected to change upon mutating the sidechain to serine, a likely explanation for the reduction in affinity is that introducing the change would distort the HA-binding site and thus decrease the affinity for HA. It is beyond the scope of the present study to determine whether defective anchoring to HA stemming from mutations in the binding site cause osteochondritis dissecans, but the availability of the ACAN-HA structure may now provide an opportunity to interpret the effect of ACAN mutations found in human patients.

### The G2 domains of ACAN does not bind to HA in spite of residue conservation in the HA- binding site

With the ACAN-HA structure determined, it was of interest to examine the conservation of HA-binding residues in the tandem Link domains found in the G2 region of ACAN. In humans, these domains share 65% amino acid identity with those found in the G1 domains (Fig. 5A). Furthermore, comparison of the Link-1 and Link-2 domains of ACAN(G1) with a model of the G2 domain generated by AlphaFold indicates a high degree of structural similarity as the two regions superimpose with a RMSD of 0.72 Å over 199 Cα atoms (Fig. 5B). Examination of the conservation of HA-binding residues indicates that 15 of 18 residues that contact HA in G1 are conserved in G2. The only changes, G219P, S298Y, and L322K in humans, correspond to residues that contact GlcNAc3, GlcNAc9, and GlcNAc7, respectively, in the crystal structure (Fig. 2C and 5B).

It was thus puzzling that a previous report suggested that the G2 segment does not bind to HA given the high degree of structural and sequence similarity between the Link-1-Link-2 and Link-3-Link-4 pairs (14). However, BLI assays conducted with purified human ACAN(G2) showed no quantifiable interaction with HA, in agreement with prior reports (Fig. 5C). Despite this result, we wondered whether the presence of the G2 domain would alter the affinity of the G1-G2 segment of ACAN for HA when compared to the G1 region only. BLI assays were thus carried out using purified ACAN(G1-G2). The K_D_ for the interaction with immobilized HA was measured at 313 nM (Fig. 5D). This affinity constant is slightly higher than the 234 nM measured for between ACAN(G1). The differences between the values of K_D_ measured for ACAN(G1) and ACAN(G1-G2) are statistically significant, and are entirely due to differences in the on rates, because there is no statistical differences between the off-rates (Fig. S6).

Furthermore, introducing alanine residues in place of R163, R217, R326, and Q334 in ACAN(G1-G2) ablated HA-binding activity, as was the case for ACAN(G1) (Fig. 3C and 5E). As such, our biochemical analyses confirm previous reports that the G2 domain of ACAN does not bind to HA. These results also indicate that the HA-binding activities of the G1 and G1-G2 regions of ACAN are similar.

To provide a rationale for the lack of HA-binding activity in G2, the changes G219P and S298Y were introduced in ACAN(G1) and the variant tested in HA-binding assays using BLI (Fig. 5F). We did not introduce the change L322K because the K647 in G2 points away from the HA-binding site and is thus unlikely to interfere with HA binding. Introduction of both the G219P and S298Y changes in ACAN(G1) leads to a 16-fold decrease in affinity for HA compared to wild type (Fig. 3A and 5F). Interestingly, the association rate constant between HA and the variant was reduced about tenfold while the dissociation rate constant is almost identical to wild type.

These findings suggest that amino acid changes in the HA-binding site contribute partly to the lack of interaction between ACAN(G2) and HA. However, additional structural alterations in G2 compared to G1 that are not currently predicted by AlphaFold may explain the absence of HA-binding affinity in this domain. These may be identified when experimental structural information on G2 becomes available.

### The HA-binding activity of ACAN is important for its integration into PNNs

In addition to being an essential component of the cartilage ECM, ACAN plays a critical role in the structure and function of PNNs, a condensed form of neuronal ECM rich in HA (19). As such, it was of interest to investigate the importance of the HA-binding activity of ACAN to its integration in PNNs. To accomplish this objective, we assessed the binding of ACAN fragments to dissociated cultures of cortical neurons. PNNs can form in such primary neuronal cultures and this model system has been used in the past to investigate the role of phosphacan in the assembly of PNNs (28, 39). Endogenous Acan localizes to nets found in cortical cultures, so another component of PNNs, the chondroitin sulfate proteoglycan Neurocan (Ncan) was used to mark their presence (25, 40). To differentiate between endogenous Acan and fragments added to cultures, regions of ACAN were expressed as V5-tagged fusion proteins and purified from the conditioned media of HEK293 cells.

We first evaluated the binding of the V5-tagged G1 and G1-G2 domains of ACAN to PNNs in primary neuronal cultures (Fig. 6). Interestingly, both fragments of ACAN bound highly and very specifically to PNNs on cultured cortical neurons as evidenced by the similar staining patterns obtained with antibodies against V5 or against the PNN component Ncan. The correlation coefficient R is above 0.95 for the binding of ACAN constructs to endogenous Ncan PNN staining (Fig. 6A). To carefully assess potential differences in binding affinity or specificity, ACAN fragments were added to cultures in three distinct amounts and concentration curves were generated (Fig. 6B). In these experiments, we determined that exogenous addition of constructs at concentrations between 0.5 nM and 50 nM represented the ideal range to study interactions of ACAN fragments with net-bearing neurons. Little binding was detected at concentrations below 0.5 nM while there was little change in PNN binding intensity but more aggregation and non-specific binding at concentrations above 50 nM (data not shown). Significant binding compared to untreated cells was measured at both 5 nM and 50 nM for both the G1 and G1-G2 fragments (Fig 6B, ACAN(G1): 5 nM p = 0.0063, 50 nM p = 0.0002; ACAN(G1-G2): 5 nM p = 0.0015, 50 nM p <0.0001). In these experiments, the association of the G1 and G1-G2 regions to net-bearing neurons was not significantly different.

**Figure 6.**
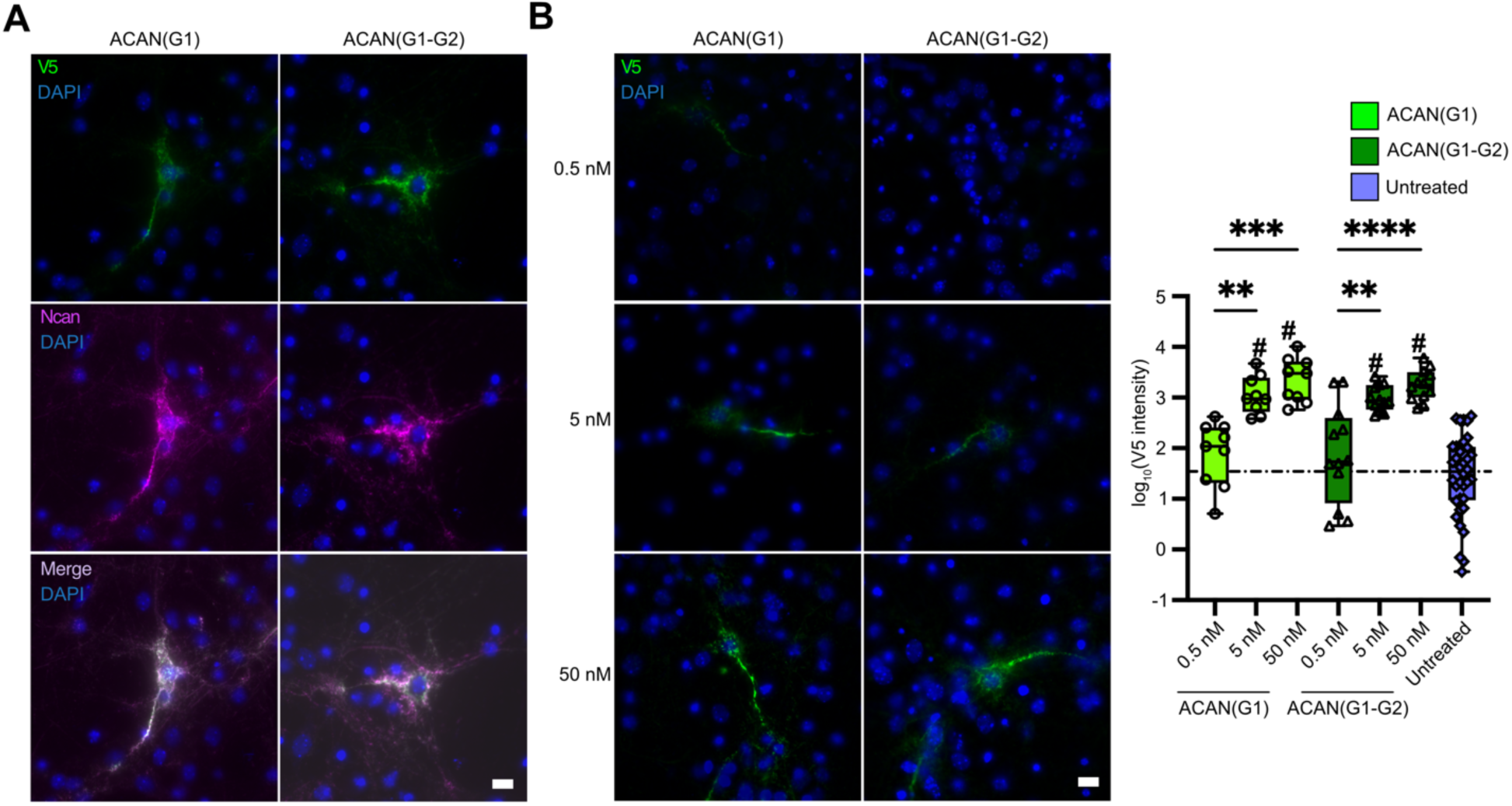
ACAN(G1) and ACAN(G1-G2) fragments bind specifically to PNNs. A. Purifed wild type ACAN(G1) and ACAN(G1-G2) [ACAN(G1)-WT and ACAN(G1-G2)-WT] tagged with V5 at 5 nM bound highly and specifically to PNNs as shown by the colocalization of V5 reactivity with PNN marker Ncan on cultured cortical neurons (V5 colocalization to Ncan reactivity, R = 0.971 ± 0.032 and 0.965 ± 0.037 for ACAN(G1) and ACAN(G1-G2) respectively by Manders’ colocalization coefficient). N = 4 independent cultures, 3 large tiled (1259 μm X 941 μm) images per culture per condition. Scale bar 10 μm. B. Binding of wild type ACAN(G1) and ACAN(G1-G2) assessed by V5 reactivity showed optimal binding between 0.5 nM and 50 nM. This range appears ideal to study the interactions of ACAN fragments with PNNs. Binding at 5-nM and 50-nM doses for both ACAN(G1) and ACAN(G1G2) was specific and intense PNN binding was observed compared to untreated control cells (Ordinary one-way ANOVA, <0.0001, F (6,92)= 3.807; Tukey’s *post hoc* testing, # indicates p < 0.0001 compared to untreated control group). Cells treated with 5-nM and 50-nM doses also showed significantly greater binding than at 0.5 nM for both ACAN(G1) (5 nM, p = 0.0063, 50 nM p = 0.0002) and ACAN(G1-G2) (5 nM, p = 0.0015, 50 nM p <0.0001) fragments. N = 4 independent cultures, 3 large tiled (1259 μm X 941 μm) images per culture per condition. Scale bar 10 μm.

We next compared the binding of the wild type and HA-binding mutant of ACAN to PNNs using both the G1 and G1-G2 regions. Since it appeared that a concentration of 50 nM provided optimal binding for wild type constructs, we utilized this concentration to assess differences in PNN binding between the wild type and HA-binding mutant constructs. The intensity of binding of the HA-binding mutants to PNNs compared to wild type constructs was significantly reduced by almost 5-fold (Fig. 7, ACAN(G1)-mutant: 24.8%, p = 0.0002, ACAN(G1-G2)-mutant: 19.6%, p = 0.0014). However, quite surprisingly, the HA-binding mutant constructs still bound specifically to PNNs, albeit with signicantly lower intensity. Importantly, as with the wild type constructs, concentrations above 50 nM did not increase PNN binding but instead lead to non-specific binding and clumping (data not show). At no concentration did the intensity of PNN binding of the HA-binding mutant constructs became equivalent to that of the wild type constructs. These data suggest that HA binding is important for ACAN integration into to PNNs but is not required for the specificity of ACAN binding to PNNs.

**Figure 7.**
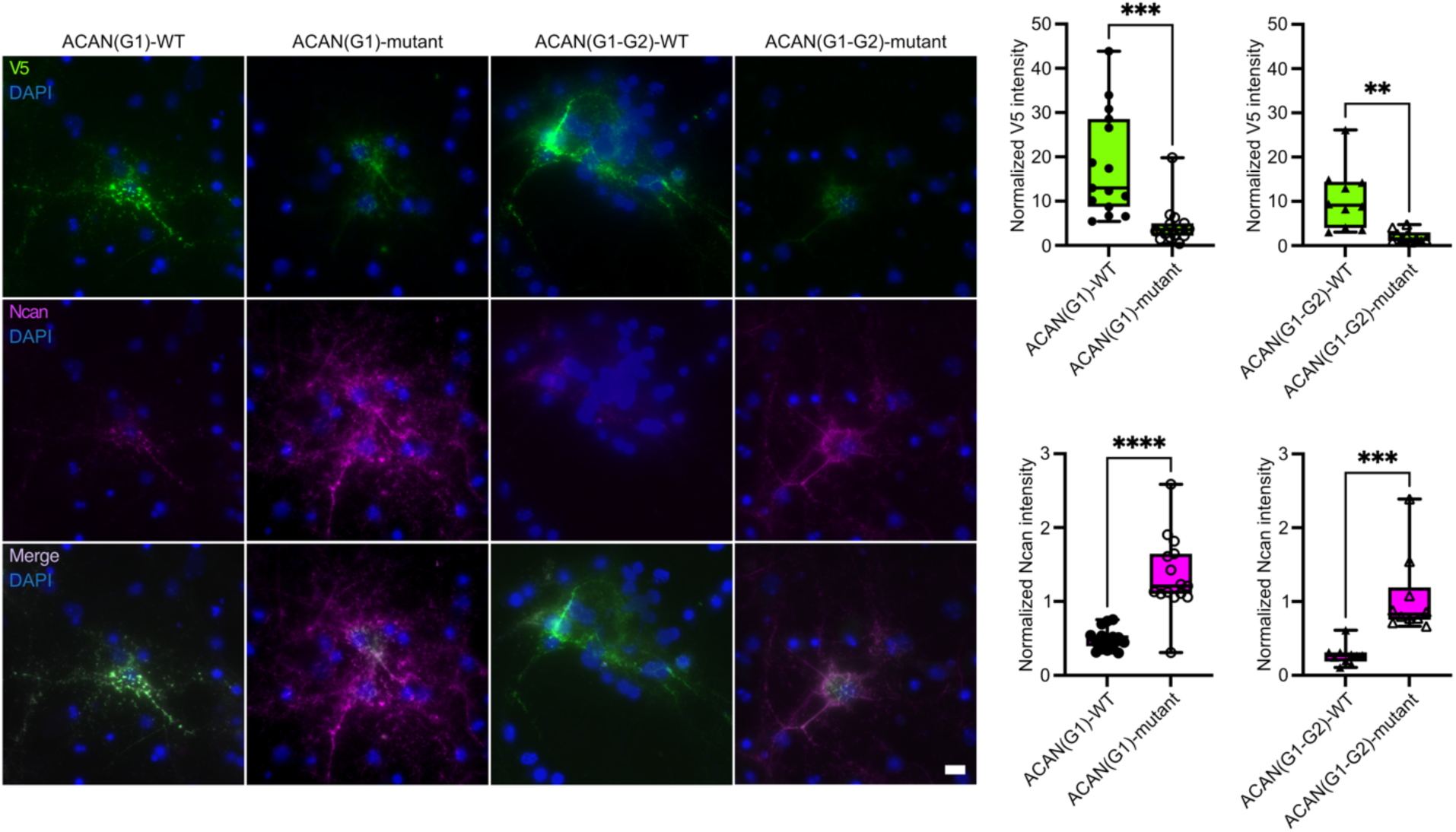
The HA-binding activity of ACAN is required for high-affinity binding, integration into PNNs, and disruption of PNN structure but is not required for specific binding to PNNs. Purified V5-tagged wild type and HA-binding mutant forms of ACAN(G1) [ACAN(G1)-WT and ACAN(G1)-mutant] and ACAN(G1-G2) [ACAN(G1-G2)-WT and ACAN(G1-G2)-mutant] were exogenously added to primary neuronal cultures at a final concentration of 50 nM. Surprisingly, both the wild type and mutant constructs localized specifically to PNNs. However, the HA-binding mutants showed significantly lower V5 staining intensity (Binding: ACAN(G1)-mutant 24.8%, p = 0.0002, ACAN(G1-G2)-mutant 19.6%, p = 0.0014). To assess the impact of these constructs on PNNs we co-labeled with the endogenous PNN marker Ncan. Data were normalized to untreated control cells (a value of 1 representing equivalent Ncan staining to untreated controls). Wild type ACAN(G1) and ACAN(G1-G2) fragments disrupt Ncan staining at significantly higher levels compared to their mutant counterparts (Ncan intensity compared to HA-binding mutant: ACAN(G1)-WT 36.16%, p < 0.0001, ACAN(G1-G2)-WT 26.38%, p = 0.0003). The mutant constructs showed no disruption of NCAN staining in PNNs compared to untreated cells. Unpaired t-test, N = 4 independent cultures, 3-5 PNNs images per culture per condition. Scale bar 10μm.

Although the role ACAN plays in the formation of PNNs is still being defined, the sum of the available data suggests that it forms a molecular bridge in PNNs by binding HA via its N-terminus and the essential PNN component TNR via its C-terminus. Therefore, we hypothesized that exogenous addition of N-terminal fragments of ACAN would displace full-length endogenous Acan, which would impair the molecular bridge to TNR and thus disrupt PNNs. To assess the impact of these constructs on PNNs we analyzed staining of the PNN component Ncan. In line with our hypothesis, the wild type ACAN(G1) and ACAN(G1-G2) proteins disrupted Ncan staining in PNNs significantly at the 50 nM concentration (Fig. 7, ACAN(G1)-WT: 36.16%, p < 0.0001, ACAN(G1-G2)-WT: 26.38% of untreated control levels, p = 0.0003). In contrast, the variants of ACAN(G1) and ACAN(G1-G2) lacking HA-binding activity did not disrupt Ncan staining in PNNs. Therefore, although HA binding is not required for specific binding of the N-terminal constructs of ACAN to PNNs, it plays an important role in maintaining the integrity of PNNs at the neuronal surface.

## DISCUSSION

PNNs were identified more than a century ago, yet it is only since the late 1990s that analyses of their molecular components and the molecular interactions that promote the assembly of nets have been undertaken. Although it is widely accepted that HA is abundant in PNNs (41, 42), the roles played by HA and protein-HA interactions in the assembly of nets are less clear cut. For example, experiments aiming to reconstitute PNNs in HEK293 cells show that expression of HAS3 along with the essential net components ACAN and HAPLN1 promotes the formation of PNN-like structures around the cells (18). HAS3 is the hyaluronan synthase most frequently expressed in PNN-bearing cells (42), yet the number of neurons bearing nets is not reduced in the hippocampus or cortex of *Has3*^-/-^ mice (43). With these somewhat conflicting considerations in mind, it was of interest to analyze the importance of the HA-binding activity of ACAN to its integration into PNNs.

To achieve this objective, the crystal structure of the N-terminal G1 domain of ACAN was determined in the presence of an HA decasaccharide, which made it possible to engineer a variant of ACAN that does not associate with HA. The abilities of the G1 and G1-G2 domains ACAN to bind to PNN-bearing neurons from mouse cortical cultures were then tested at distinct concentrations. In these experiments, the G1 and G1-G2 regions behaved in essentially identical fashion: they could both bind to net-bearing neurons and inhibit the integration of Ncan in PNNs. What was perhaps unexpected is that the mutant forms of ACAN(G1) and ACAN(G1- G2) that do not bind HA in the BLI binding assays retained some reduced but significant association with net-bearing neurons compared to wild type proteins. The simplest conclusion from these results is that ACAN can assemble into nets through two distinct activities: its binding to HA as well as interaction with another net component, perhaps through protein-protein interactions. In the latter case, one may speculate that HAPLN1, a known ACAN-binding protein, would promote the integration of ACAN into nets through interactions between their Ig domains (15). HAPLN1, in turn, could be immobilized to the neuronal surface through interactions with HA synthesized by a hyaluronan synthase. Yet, even though the presence of PNNs is reduced in mice lacking Hapln1, some residual Acan still localizes to neurons bearing attenuated nets (17). Obviously, this result could be interpreted as Acan immobilizing on neurons owing to its interactions with HA, However, at this stage, it is not possible to exclude that a yet undescribed protein may function as a receptor for ACAN on the surface of net-bearing neurons.

The potential ability of ACAN to assemble into PNNs because of protein-protein as well as protein-HA interactions would also explain a somewhat counter-intuitive result. Net-like structures that include Hapln1 and Tnr persist in dissociated cortical cultures of cartilage matrix deficiency (cmd) mice (20). In cmd mice, a deletion of 7 base pairs in the gene encoding Acan results in the expression of a truncated protein that includes the N-terminal Ig domain and part of the Link-1 module (9). Based on the results reported here, this truncated protein would be unable to bind HA. Homozygous mice carrying this mutation die soon after birth, preventing investigation of PNNs in their neural tissues. Apart from considering possible compensatory mechanisms to mitigate defective Acan secretion (44), the presence of PNN-like structures in dissociated cortical cultures from cmd mice is difficult to explain except if the truncated Acan protein expressed by these cells is sufficient to induce condensation of the matrix around neurons, even if it is unable to bind HA. Thus, we hypothesize that the N-terminus of Acan, and perhaps more specifically its N-terminal Ig domain mediates protein-protein interactions that promote the formation of PNNs.

Another interesting question centers around the role of ACAN in PNNs. When PNNs were reconstituted on the surface of HEK293 cells, the expression of ACAN, HAPLN1, and HAS3 was necessary to obtain a condensed matrix around the cells (18). In contrast, the expression of HAPLN1 and HAS3 only resulted in a diffuse matrix. The condensed aspect of the matrix could stem from more significant cross-linking of HA chains through the combined interaction of ACAN with HA and HAPLN1. Viewed from that angle, it is troubling that the G2 domain of ACAN does not appear to bind HA in spite of the resemblance between its Link modules and those found in G1 because it could provide some of the additional cross-links needed to condense the matrix around neurons. Perhaps the G2 fragment associates with HA at concentrations well beyond those tested in our BLI assays, but these could be attained in the context of PNNs where the local respective concentrations of ACAN and HA are presumably much higher than in the assays performed here. Another possibility is that the G2 region binds to another protein, which induces changes in the structure of G2 to promote HA binding.

However, in the context of dissociated neuronal cultures, the G2 domain of ACAN did not bind to PNNs (data not shown). Thus, at this stage, we remain unable to ascribe a role to the G2 region, and additional experiments focused on identifying its binding partners are needed. Finally, a structural argument could be made that it is only the binding of HA to ACAN(G1) that promotes condensation of the matrix. Indeed, the structure reported here indicates that the conformation of HA is altered when bound to ACAN (Fig. S3). This change in conformation in the bound HA may induce a local tension in the HA strand, which repeated over the length of several crisscrossing HA chains might trigger condensation of the matrix. These possible explanations for the condensation of the matrix around neurons are not mutually exclusive. As such, future characterization of the molecular interactions mediated by the distinct domains of ACAN will provide additional keys to understand why is such an essential component of nets.

## EXPERIMENTAL PROCEDURES

### Cloning

A cDNA encoding the G1 region of human ACAN (Uniprot ID P16112, amino acids 21 – 351) was synthesized by Genscript and cloned into a derivative of the pLex2 vector (45). This vector directs the expression of a signal peptide from bovine serum albumin followed by an hexahistidine tag, a human rhinovirus 3C protease site and ACAN(G1). A cDNA fragment encoding human ACAN(G2) or ACAN(G1) with the mutations R163A, R217A, R326A, and Q334A were generated by Integrated DNA Technology and cloned into the pLex2 derivative as described above. Fragments encoding ACAN(G1) with the mutations D200E, G319S, and G219P + S298Y were synthesized by Twist Bioscience and also cloned into the same expression plasmid.

A similar strategy was used to express the G1-G2 region of human ACAN. cDNA fragments encoding amino acids 21 – 676 of wild type human ACAN or human ACAN including the changes R163A, R217A, R326A, and Q334A were synthesized by Genscript and cloned into a derivative of the pLex2 vector. In this case, however, this vector directs the expression of a signal peptide from human interleukin-2 followed by an hexahistidine tag, a human rhinovirus 3C protease site, ACAN(G1-G2), and a C-terminal V5 tag. Fragments encoding ACAN regions used in neuronal assays shown in Figures 6 and 7 were all cloned into this vector. All plasmid constructs were verified by DNA sequencing.

### Protein expression and purification

HEK293T/17 cells (ATCC # CRL-11268) used for transfection were grown in suspension in FreeStyle F17 media (ThermoFisher Scientific) supplemented with 8 mM L-Glutamine and 1% (v/v) ultra-low IgG fetal bovine serum (Gibco). Cells were maintained in vented polycarbonate flasks shaken at 135 rpm and kept at 37 °C and 8% CO2 (46). The day before transfection, cells were seeded at 8 x 10^5^ cells/ml in vented polycarbonate shaker flasks. The day of the transfection, cells were counted and diluted 1 x 10^6^ cells/ml right before adding a solution of DNA- polyethyleneimine complex. For transfecting a 100 mL culture at 1 x 10^6^ cells/ml, this solution includes 100 µg of plasmid mixed with 300 µl of a 1 mg/ml solution of polyethyleneimine (PEI MAX, Polysciences) in 4 mL of OptiPRO SFM media (ThermoFisher Scientific). The solution of DNA-polyethyleneimine was incubated for 10 – 30 minutes at room temperature before being added to the cells. The cells were maintained at 37 °C for ∼12 – 16 hours then switched to a temperature of 33 °C. Cultures were harvested 5 to 6 days after. When appropriate, kifunensine (Carbosynth) was added to a final concentration of 7.5 µM to cells the day of the transfection to generate proteins with paucimannose N-linked carbohydrates that can be removed by endoglycosidases prior to initiating crystallization trials (47).

Transfected cells and conditioned media were separated by centrifugation and conditioned media was supplemented with solutions of 400 mM NaPO4 pH 7.5 and 5 M NaCl to final concentrations of 40 mM and 500 mM, respectively. The media was then applied to a 5-mL HisTrap excel column (Cytiva) equilibrated in 500 mM NaCl, 50 mM NaPO4 pH 7.5. The bound protein was eluted with a gradient to 500 mM imidazole over 15 column volumes. Rhinovirus 3C protease or a mixture of human rhinovirus 3C protease, endoglycosidase H, and endoglycosidase F were added to the purified protein and dialyzed overnight at room temperature against 250 mM NaCl, 50 mM NaPO4, pH 7.5, 40 mM imidazole, and passed over a 5-mL His-Trap column (Cytiva). The flow through was kept. Subsequent purification steps involved ion exchange on a 5- mL HiTrap Q HP column (Cytiva) equilibrated in 20 mM Tris-HCl pH 8.0 followed by gel filtration chromatography on a Superdex 75 26/600 column (Cytiva) equilibrated in 150 mM NaCl and 20 mM Na-HEPES pH 7.5.

For PNN-integration assays, constructs expressing V5-tagged ACAN fragments were transfected into adherent HEK293 cells using a PEI transfection method. Briefly, for each 100 mm plate of HEK293 cells, 10 μg of DNA construct was added to 1 mL of serum free DMEM (Thermo Fisher Scientific). A 21-μL aliquot of PEI at 1 mg/mL was added to the DNA solution and incubated at room temperature for 15 minutes. The transfection mix was added to a 100-mm plate at ∼ 80% confluency. Number of plates and volumes were scaled as required. Cells were switched to serum-free Opti-MEM media (Thermo Fisher Scientific) 24 hours after transfection. Conditioned media from transfected HEK293 cells were collected 48 hours post transfection and concentrated using a 30,000 MWCO concentrators (AmiconUltra, EMD Millipore). Protein of interest from the concentrated media was purified using Cobalt Spin Columns (Thermo Fisher Scientific). Samples were mixed with wash buffer (50 mM NaPO4, 300 mM NaCl, 10 mM imidazole; pH 7.4) and applied to the column. Columns containing samples were incubated for at least 30 minutes at 4°C in an orbital or end over end shaker. Columns were washed 3 times with wash buffer and protein was eluted using 50 mM NaPO4, 300 mM NaCl, 150 mM imidazole; pH 7.4.

For addition to neurons, eluted protein from above was desalted using Zeba spin desalting columns (Thermo Fisher Scientific) as per the manufacturers protocol and concentrated using Pierce protein concentrators (Thermo Fisher Scientific). Samples were diluted to 1 µM in Neurobasal media (Thermo Fisher Scientific) and added to cells as required.

### Crystallization, structure determination, and structural analyses

Crystals were grown at 20 °C by hanging drop vapor diffusion. For crystallization of ACAN(G1), a 1-µl aliquot of deglycosylated protein at 7.4 mg/ml in 10 mM HEPES pH 7.5, 75 mM NaCl was mixed with 1 µl of 1% (w/v) PEG 3,350, 100 mM Bis-Tris, pH 5.5, and 600 mM (NH4)2SO4. Crystals were frozen in mother liquor supplemented with 30% (v/v) glycerol. For crystallization of the ACAN(G1)-HA complex, a 1-µl aliquot of deglycosylated protein at 7.4 mg/ml (∼200 µM) in 10 mM HEPES pH 7.5, 75 mM NaCl was mixed with 1 µl of a 1 mM solution of HA decasaccharide in H2O (HA_10_, Iduron, distributed by Galen Laboratory Supplies) and 1 µl of 15% (w/v) PEG 10,000, 50 mM Bis-Tris propane, pH 9.0, and 200 mM NH4C2H3O2. Crystals were frozen in mother liquor supplemented with 20% (v/v) PEG200 and 50 µM of HA_10_.

X-ray diffraction data were collected on beamline 22-ID of the Advanced Photon Source at Argonne National Laboratory. Diffraction data were processed using HKL2000 (48). Structures were determined by molecular replacement in PHASER as implemented in PHENIX (49, 50) using a model of human ACAN(G1) generated with AlphaFold2 (32). The final models were obtained after several rounds of manual rebuilding in COOT (51) and refinement in PHENIX. These models were validated using the RSCB PDB validation server. List of interacting residues and buried surface area were generated with Chimera X (52). RMSD values between superimposed structures were calculated with GESAMT (53) implemented in CCP4 (54). For structural comparisons, structures were superimposed using the least-square or secondary structure matching options in COOT. All structural representations were generated using ChimeraX.

### HA-binding assays using biolayer interferometry

Interactions between fragments of human ACAN and HA were quantified at room temperature in 150 mM NaCl, 10 mM Na-HEPES pH 7.5, 1 mg/mL BSA, 0.02 %(v/v) Tween-20 using an Octet K2 system (Sartorius). A 20-kDa fragment of HA with a single biotin molecule per chain (Creative PEGWorks, 250 nM) was immobilized onto streptavidin tips (Sartorius). These tips were then incubated with purified ACAN proteins at a series of concentrations for 180 s during the association phase. The tips were then incubated in buffer only during the dissociation phase for 180 s. The signal was corrected by subtracting the background measured for the buffer only. Experimental data were analyzed using a 1:1 binding model as implemented in the Octet Data Analysis software (version 11.0.0.4). For visualization, sensorgrams were plotted using Prism 10 (GraphPad Software). The results are reported as the average of at least four replicates. These include at least two independent measurements for each of at least two distinct batches of proteins. Additional information about individual experiments used in affinity calculations are listed in Table S1.

### Sequence alignments

Amino acid sequences for aggrecan for human (Uniprot ID P16112), mouse (Uniprot ID Q61282), chicken (Uniprot ID P07898), and zebrafish (Uniprot ID F1QDA1 for Acana and Uniprot ID A0A8M9P6I2 for Acanb) were aligned using Clustal Omega available from the European Bioinformatics Institute website (55).

### Animals

For primary cortical neuronal cultures, timed pregnant CD-1 WT mice were purchased from Charles River Laboratories (Wilmington, MA). All experiments followed the protocols approved by the Institutional Animal Care and Use Committee of SUNY- Upstate Medical University.

### Primary cortical cultures

Primary cortical neuronal cultures were prepared as described previously (20, 56). Briefly, cortices from embryonic day (E) 16 CD-1 WT embryos were dissected out and digested with 0.25 % trypsin-EDTA (ThermoFisher Scientific) for 25 minutes. The tissue ball after trypsin digestion was treated with RNA-ase free DNA-ase (Promega) for 6 minutes and passed through a 70 µm cell strainer (Falcon). Cells were centrifuged to remove any residual DNA-ase and resuspended in Neurobasal medium supplemented with B27, GlutaMAX and penicillin-streptomycin (ThermoFisher Scientific). Cultures were plated at a density of 2.1 x 10^6^ cells/mL on glass coverslips precoated with poly-D-lysine (100 µg/ml, Sigma-Aldrich) and laminin (50 µg/ml, ThermoFisher Scientific). Cells were treated with 5 µM cytosine arabinoside (AraC, Sigma-Aldrich) from 1 – 3 days *in vitro* (DIV) to eliminate glia. Culture media was replaced at 3 DIV after AraC treatment, followed by a half media change at 6 DIV. For PNN integration experiments, purified ACAN fragments tagged with V5 (see above) were added to cells at 6 DIV as required. Cells were maintained at 37 °C, 5% CO2 until fixation at 9 DIV.

### Immunocytochemistry

Primary cortical cultures plated on coverslips were fixed in cold 4% phosphate-buffered paraformaldehyde (PFA) with 0.01% glutaraldehyde, pH 7.4 at 9 DIV. Afterwards, the cells were blocked in screening medium (DMEM, 10% (v/v) FBS, 0.2% (w/v) sodium azide) for 1 h, before adding primary antibodies overnight at 4 °C (rabbit anti-V5: invitrogen MA5-32053; sheep anti- Ncan: biotechne AF5800). The next day, Alexa Fluor– conjugated secondary antibodies (ThermoFisher Scientific) in screening medium were added to the cells for 2 h before mounting the coverslips with ProLong Antifade Kit (ThermoFisher Scientific). Cell nuclei were visualized with Hoechst solution (ThermoFisher Scientific) diluted in PBS. For quantification of probe binding, large tiled images (1259 µm X 941 µm) were taken using a 20x objective (NA=0.8). Images were converted to 8-bit and background correction was carried out by subtracting modal value from images. Intensity of binding was determined using the measure function in ImageJ. For Ncan disruption experiments, individual PNNs were imaged using a 40x oil objective (NA=1.4). Background subtraction was carried out using the rolling ball background correction function in ImageJ. The intensity of Ncan in PNNs was determined in ImageJ using the measure function.

## DATA AVAILABILITY

The atomic coordinates and structure factors (codes 9DFT, 9DFF) have been deposited in the Protein Data Bank (https://www.rcsb.org). All other data are contained within the article and Supporting Information.

## SUPPORTING INFORMATION

This article contains supporting information.

## ACKNOWLEDGEMENTS

We thank Dr. Xiaolan Yao for use of her Octet K2 system.

## FUNDING AND ADDITIONAL INFORMATION

Research reported in this manuscript was supported by the National Institute of General Medical Sciences under award number R01 GM143757 (R.T.M. and S.B.). The content is solely the responsibility of the authors and does not necessarily represent the official views of the National Institutes of Health. Data were collected at Southeast Regional Collaborative Access Team (SER-CAT) 22-ID or 22-BM beamlines at the Advanced Photon Source, Argonne National Laboratory. SER-CAT is supported by its member institutions (see www.ser-cat.org/members.html), and equipment grants (S10 RR25528 and S10 RR028976) from the National Institutes of Health. Use of the Advanced Photon Source was supported by the U. S. Department of Energy, Office of Science, Office of Basic Energy Sciences, under Contract No. W-31-109-Eng-38. Figures were prepared using UCSF ChimeraX, developed by the Resource for Biocomputing, Visualization, and Informatics at the University of California, San Francisco, with support from National Institutes of Health R01-GM129325 and the Office of Cyber Infrastructure and Computational Biology, National Institute of Allergy and Infectious Diseases.

## AUTHOR CONTRIBUTIONS

Conceptualization: R.T.M. and S.B. Data curation: R.T.M. and S.B.

Formal analysis: A.S., R.T.M., and S.B. Funding acquisition: R.T.M. and S.B.

Investigation: M.Y.O., L.B.E, A.S., G.N., S. M. M., R.T.M., and S.B.

Methodology: R.T.M. and S.B.

Project administration: R.T.M. and S.B. Supervision: R.T.M. and S.B.

Writing – original draft: R.T.M. and S.B. Writing – review and editing: R.T.M. and S.B.

## ABBREVIATIONS

ACAN,: aggrecan;
ANOVA,: analysis of variance;
BLI,: biolayer interferometry;
cDNA,: complementary DNA;
CNS,: central nervous system;
DAPI,: 4′,6-diamidino-2-phenylindole;
DIV,: day in vitro; E, embryonic day;
ECM,: extracellular matrix;
PDB,: Protein Data Bank;
PNN,: perineuronal net;
RMSD,: root mean square deviation

---

**Figure S1.**
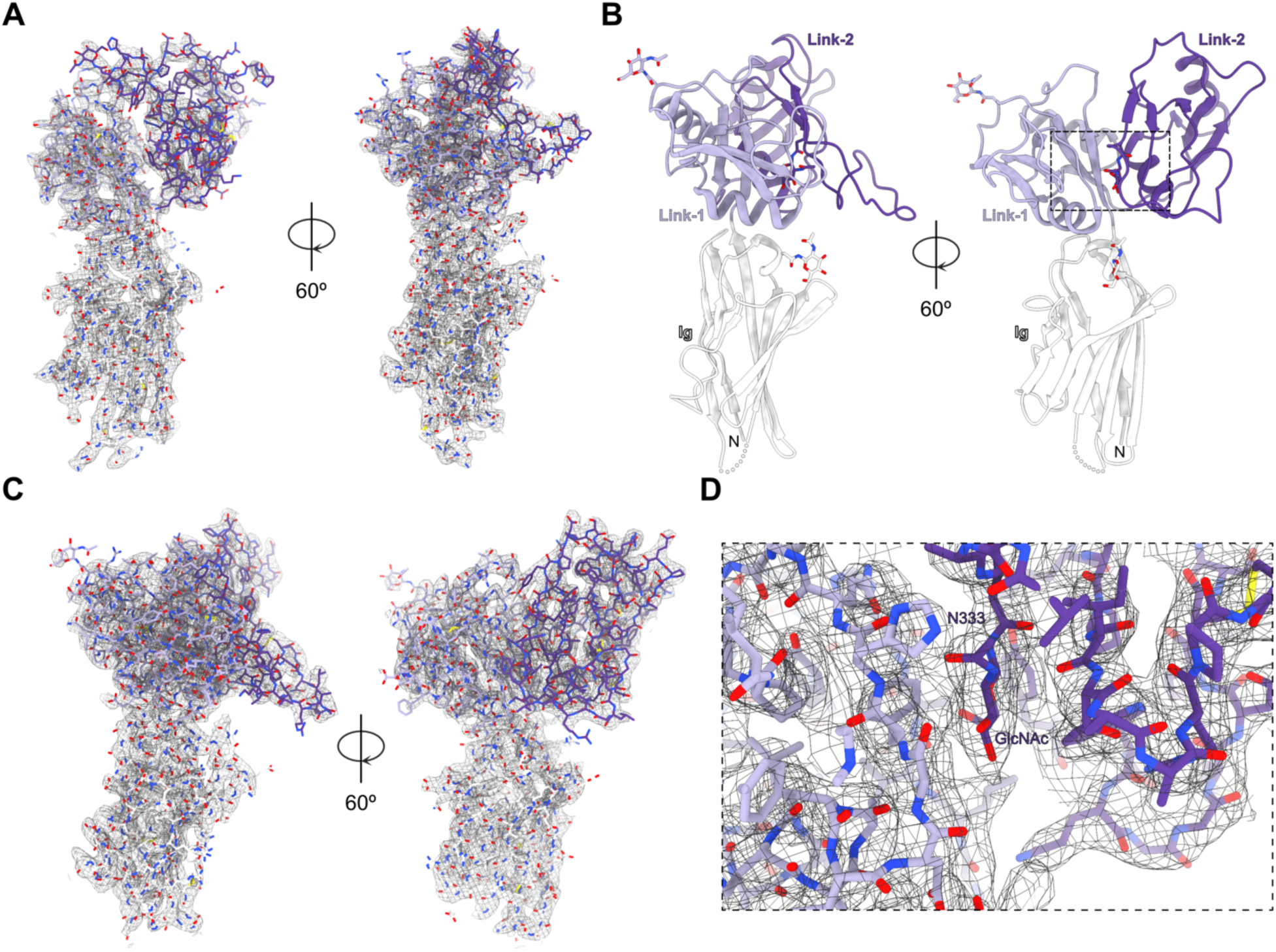
Electron density maps for the two chains of ACAN found in the asymmetric unit of the ACAN(G1) crystal structure. A. 2mFo-DFc electron density map for chain B of ACAN(G1) at 3.5 Å shown in Fig. 1 is contoured at 1.2 σ. ACAN(G1) is shown as sticks and the Ig, Link-1, and Link-2 domains are colored white, lilac, and violet, respectively. The two views are in the same orientation as the views shown in Fig. 1B. B. The G1 region of chain A of human ACAN is shown in a ribbon diagram in two distinct orientations related by a 60° rotation. The Ig, Link-1, and Link-2 domains in chain B of ACAN are colored white, lilac, and violet, respectively. The letters N and C indicate the N- and C-termini, respectively. Asparagine-linked N-acetylglucosamine residues are shown as sticks along with the asparagine side chain. A disordered region in the the Ig domain is shown as a dotted line. An area boxed by a dotted line shows the GlcNAc residue on N333. This region is shown in more details in panel D. C. 2mFo-DFc electron density map for chain A of ACAN(G1) at 3.5 Å shown in panel B is contoured at 1.2 σ. ACAN(G1) is shown as sticks and the Ig, Link-1, and Link-2 domains are colored white, lilac, and violet, respectively. The two views are in the same orientation as the views shown in panel B. D. 2mFo-DFc electron density map for region surrounding N333 in chain A of ACAN(G1) at at 3.5 Å shown in panel B is contoured at 1.2 σ. ACAN, aggrecan; GlcNAc, *N*-acetylglucosamine; Ig, immunoglobulin;

**Figure S2.**
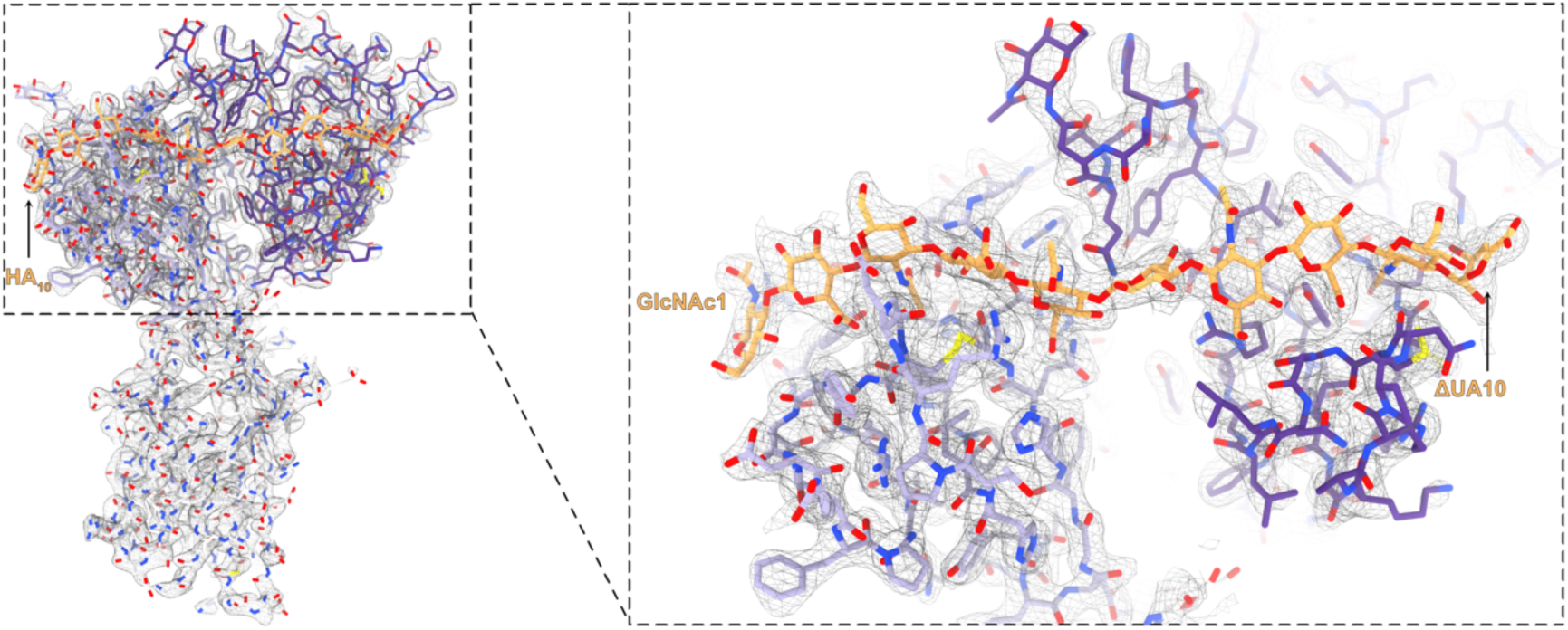
Electron density map for the ACAN-HA_10_ complex. Representative electron density maps at the interface. The ACAN-HA_10_ complex is shown here in stick representation along with a 2mFo-DFc electron density map contoured at 1.2 σ. The Ig, Link-1, and Link-2 domains of ACAN are colored white, lilac, and violet, respectively. The bound glycosaminoglycan is in orange. The boxed inset shows a close-up view of the HA-binding site in ACAN. ACAN, aggrecan; GlcNAc, *N*-acetylglucosamine; HA, hyaluronan; ΔUA, 4-deoxy-alpha-L-threo-hex-4-enopyranuronic acid.

**Figure S3.**
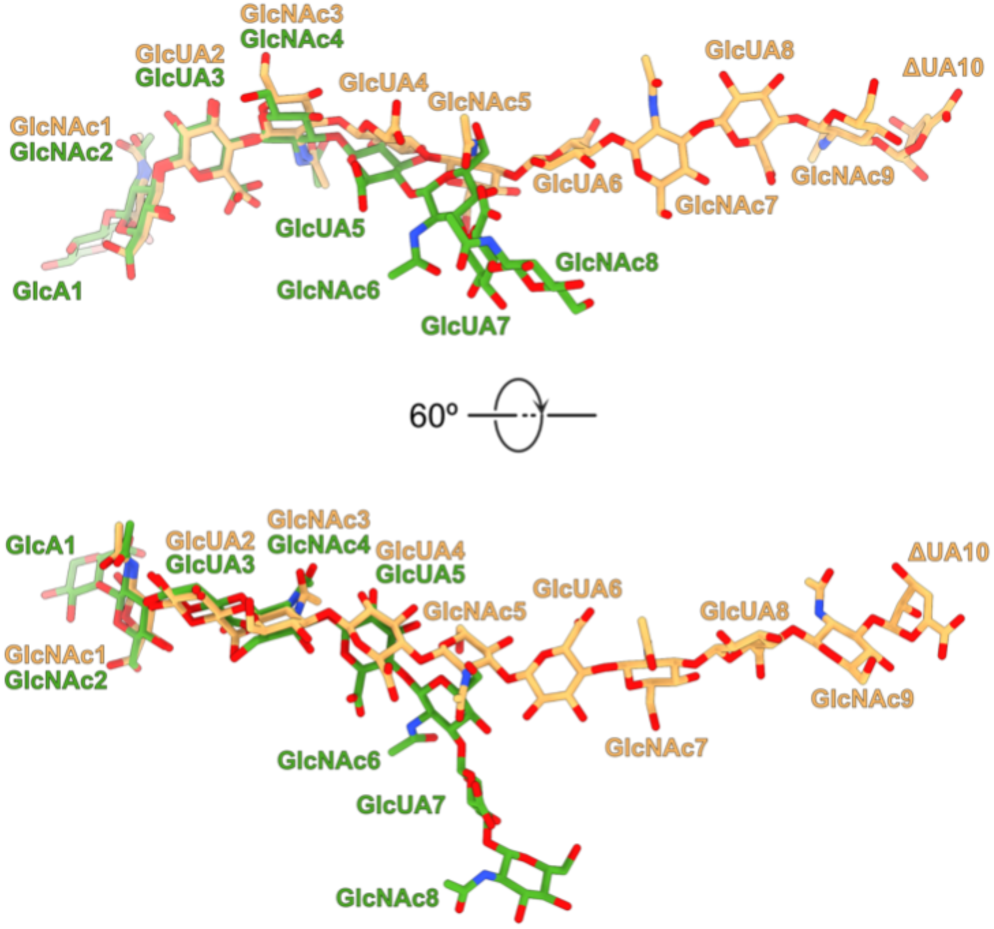
Comparison of HA oligosaccharides from powdered diffraction and in the crystal structure of the ACAN-HA complex. The HA oligosaccharide bound to ACAN is shown here as orange sticks. A structure of an HA octasaccharide determined in the absence of bound protein (PDB ID 3HYA) is superimposed onto HA_10_ using residues GlcNAc1 of HA_10_ and GlcNAc2 of “free” HA. In the top view, the HA_10_ oligosaccharide is in the same orientation as the one shown in Fig. 2A. ACAN, aggrecan; GlcNAc, *N*-acetylglucosamine; GlcA, alpha-D-glucopyranuronic acid; GlcUA, beta-D-glucopyranuronic acid; HA, hyaluronan; ΔUA, 4-deoxy-alpha-L-threo-hex-4- enopyranuronic acid.

**Figure S4.**
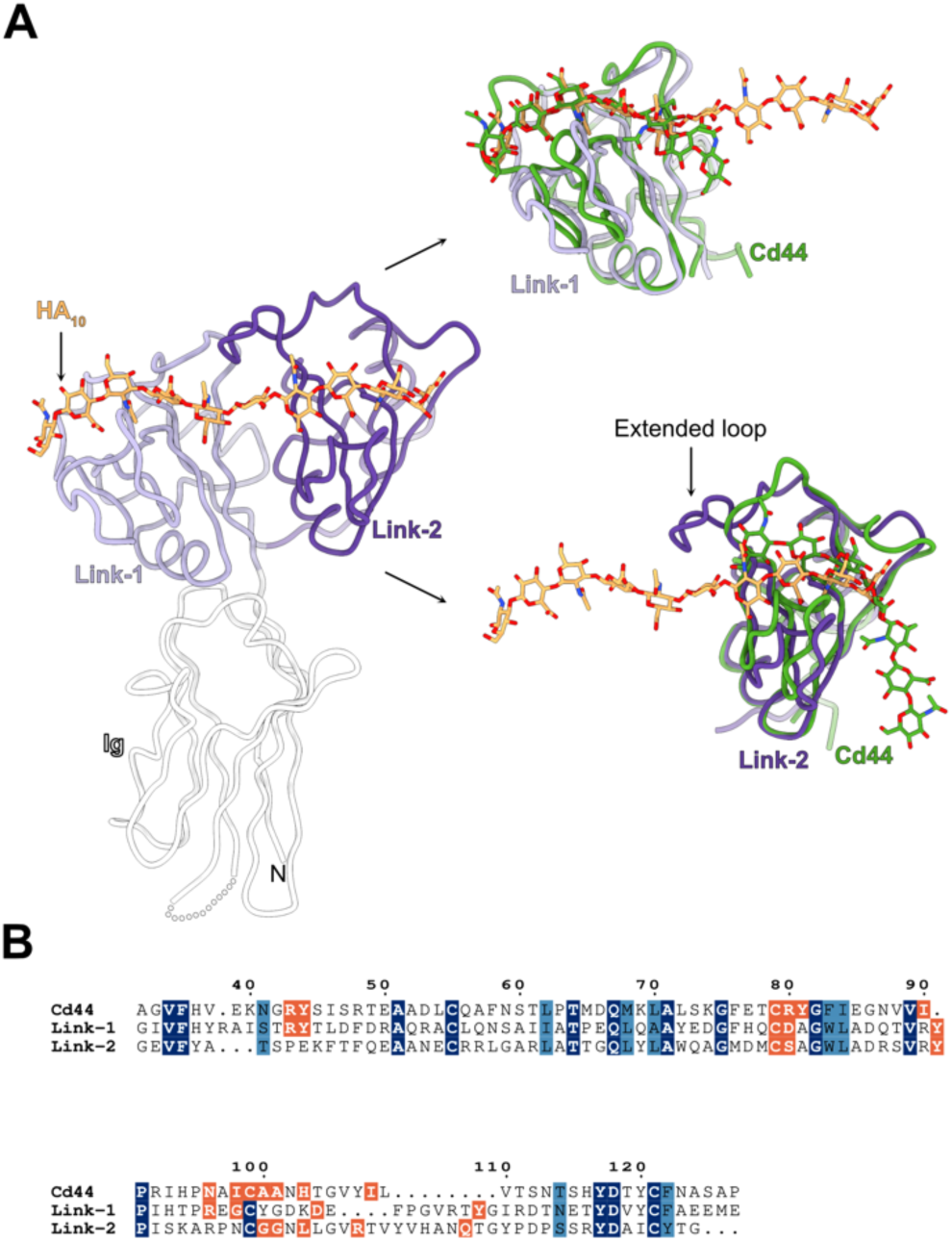
Comparison of Cd44-HA complex with the HA_10_-bound Link-1 and Link-2 domains of human ACAN. A. The bound HA_10_ oligosaccharide in ACAN occupies grooves in the Link-1 and Link-2 modules of ACAN that match the site where HA binds to the Link module in mouse Cd44 (PDB ID 2JCR). Proteins are shown in coil representation while the bound oligosaccharides are shown as sticks. The Link module of mouse Cd44 was superimposed onto the two Link modules of ACAN and colored green. The Ig, Link-1, and Link-2 domains in ACAN are colored white, lilac, and violet, respectively. The HA oligosaccharide bound to Cd44 is also colored green. A loop extends from Link-2 to bind to HA. This loop is absent in Cd44 and corresponds to an insertion of 8 amino acids between residue 109 and 110 of Cd44. B. Amino acid conservation between the Link module of mouse Cd44 and the Link-1 and Link-2 modules of human ACAN. The numbering corresponds to mouse Cd44 (Uniprot ID# P15379-14). Identical residues are shaded in navy while similar residues are shaded in blue. Residues that interact with HA are shaded orange. ACAN, aggrecan; HA, hyaluronan; Ig, immunoglobulin.

**Figure S5.**
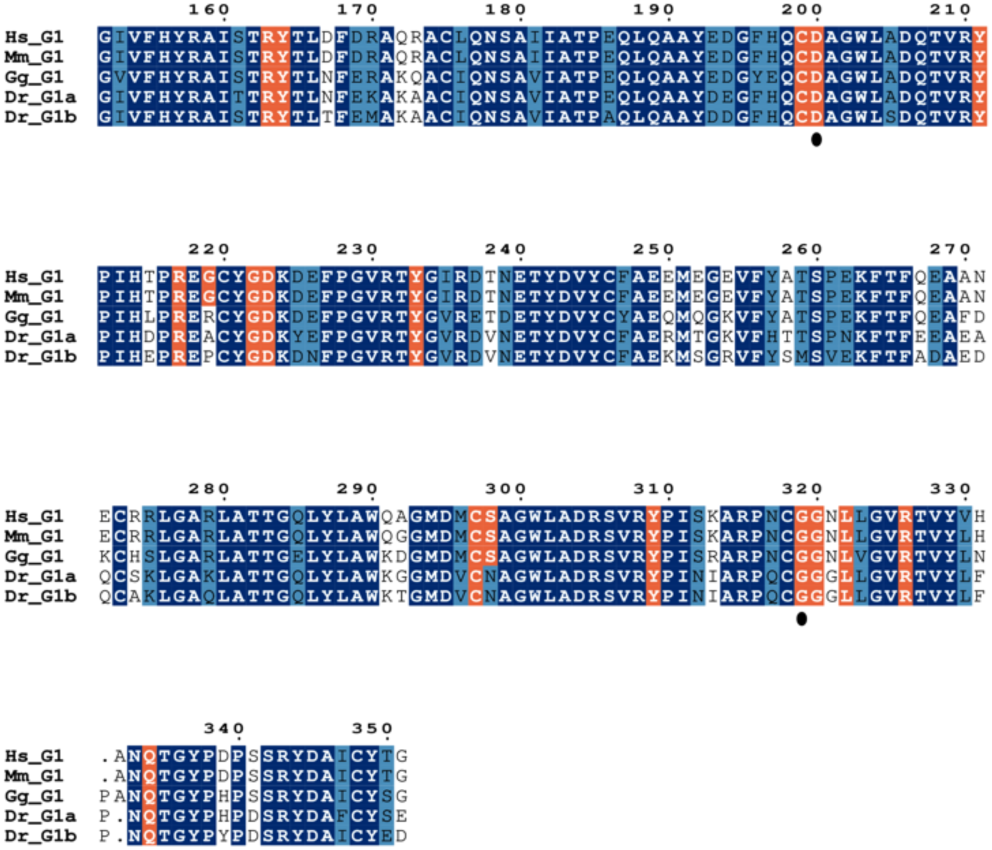
Sequence alignment of tandem Link modules found in the G1 domains of selected ACAN proteins. Amino acid conservation between the tandem Link modules of human, mouse, chicken, and zebrafish ACAN. Two genes, *acana* and *acanb*, encode homologs of human ACAN in zebrafish. The numbering corresponds to human ACAN. Identical residues are shaded in navy while similar residues are shaded in blue. Residues that interact with HA in human ACAN are shaded orange. They are highlighted in the same color in other sequences to indicate residue conservation. Black dots under D200 and G319 indicate residues that have been identified in patients suffering from osteochondritis dissecans and reported to the ClinVar database. Hs, *Homo sapiens*; Mm, *Mus musculus*; Gg, *Gallus gallus*, Dr, *Danio rerio*.

**Figure S6.**
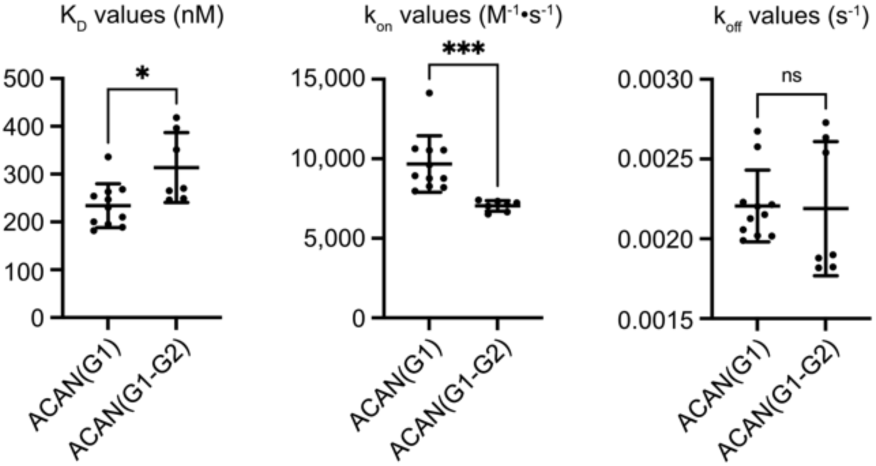
Statistical analyses of K_D_, k_on_, and k_off_ values measured for ACAN(G1) and ACAN(G1-G2). The differences between the values of K_D_, k_on_, and k_off_ for the interactions of ACAN(G1) (11 independent experiments using 3 distinct protein batches) and ACAN(G1-G2) (7 independent experiments using 3 distinct protein batches) and biotinylated HA were analyzed using a Welch t-test. The differences between the K_D_ values were significant (p = 0.0299), as were those between the k_on_ values (p = 0.0006). The values for the off-rates were not statitiscally significant.

